# Evolved microbial diversity enables combinatoric biosensing in complex environments

**DOI:** 10.1101/2025.03.24.645055

**Authors:** Alyssa Jasmine Chiang, Nicholas Csicsery, Richard O’Laughlin, Leo Baumgart, Elizabeth Stasiowski, Phuc Nguyen, Austin Doughty, Myra Ashraf, Raegan Mink, Elina C. Olson, Michael Ferry, Adam M. Feist, Nan Hao, Karsten Zengler, Jeff Hasty

## Abstract

Whole-cell biosensors (WCBs) offer rapid, cost-effective monitoring of environmental contamination and human disease. Current WCB efforts to optimize detection of single target analytes under laboratory conditions have achieved vastly improved performance, setting the stage for WCB deployment in complex environments. We propose a framework that leverages the cross-specificity of single-target WCBs to quantify multiple targets using supervised machine learning. Specifically, we engineer six sensors for heavy metal contaminants in laboratory *E. coli*. We then evolve the strain to generate five chassis with improved growth in seawater conditions. We transform the chassis with the sensors, creating a set of 30 variants. The variant dynamic responses are characterized with microfluidics, revealing significant diversity. Leveraging this diversity, we construct a consortium to combinatorically quantify multiple analytes with machine learning, outperforming single-target biosensors in over 90% of our test samples. These results form a generalizable framework that facilitates WCB translation toward settings beyond the laboratory.

## Introduction

Industrial and agricultural activities have resulted in buildup of toxic byproducts and waste, such as heavy metals, in marine environments^1–3^. As they accumulate, heavy metals harm biodiversity and present public health risks for local populations through consumption and exposure, particularly in developing countries, leading to devastating neurological and physiological damage^1,4–6^. As a result, cost-effective, rapid biomonitoring solutions that are robust to high salinity conditions are essential to facilitating pollutant mitigation and reduction efforts.

Whole-cell biosensors (WCBs) have gained traction in target analyte detection, informing biological solutions in human and environmental health^7–10^, agriculture^11^, or food safety^12^. Though chemical or physical detection methods exist for many analytes—such as mass spectrometry or high-performance liquid chromatography^13^—these assays can be expensive, time-consuming, and salinity-sensitive, while also requiring experienced personnel or complex equipment^14–16^. Whole-cell biosensors, on the other hand, not only offer affordable, versatile, rapid, and simple readouts to address these needs, but with the rise of greater precision in synthetic biology and genetic engineering^17–19^, can also be linked to real-time remedial actions such as therapeutic payload release^20,21^ or toxin neutralization^22^.

WCB development has focused largely on engineering and tuning strains for detecting single targets within a controlled laboratory environment^23^. These efforts have resulted in techniques for the detection of compounds including heavy metals^7,9,14,24,25^, caffeine^26^, aromatics^27,28^, and waterborne pathogens^29,30^. WCBs have employed a variety of techniques, including DNA-^31,32^, RNA-^31,33^, transcription factor-based^7,9,26,34–36^, and optical approaches^37,38^. More recently, evolutionary and rational engineering approaches have expanded the repertoire of possible targets for WCBs^27,28,33,36,39^. These advances have vastly improved WCB capabilities, opening the door to WCB field potential^40,41^. Currently, single-target biosensors suffer from crosstalk, preventing the simultaneous detection of multiple biomarkers in a sample^29,42^. Furthermore, WCBs characterized under laboratory settings face difficulties in survival, long-term stability, and signal interference under environmental stressors^17,43^, including temperature extremes, pH, and osmolarity, as well as exposure to toxins and oxidative compounds, limiting their deployability in complex environments^44^.

To close this gap, we shift our attention to sensing multiple target analytes in a mixed sample under high salinity seawater conditions. We engineer six transcription factor-based sensors for five heavy metals of concern in polluted seawater samples^1,3^. Then, we construct a consortium of WCBs by harnessing adaptive laboratory evolution (ALE) with the well-characterized *E. coli* K-12 MG1655 laboratory strain to efficiently generate a library of diverse biosensor signals, circumventing environmental fitness defects and the need to characterize new chassis without rational engineering that would require *de novo* information about microbe stress tolerance. To more fully capture the library’s phenotypic diversity, we develop a versatile microfluidic screening platform that can elucidate WCB dynamic behavior in continuous culture^7,45,46^. Our modular design allows parallel screening of multiple strains or conditions in a single experiment. Exploiting our WCB diversity, we expose the consortium to approximately 500 combinations of the five heavy metals in seawater to train a supervised learning regression model to quantify the metals by using the collective consortium response, analogous to a fingerprint or barcode. Instead of relying on single-target WCB responses which can be confounded by signal crosstalk, the consortium outperforms single-target biosensors in predicting heavy metal concentrations.

A microbial consortium paradigm of WCBs bridges the advancements in biosensor engineering to field deployability by shifting focus from optimization of single-target WCBs toward maximization of WCB data. The consortium-based biosensor can be transposed to a myriad of diagnostic applications that require quantification of multiple biomarkers under stressful environmental conditions.

## Results

### Engineered transcription factor-based constructs in E. coli detect heavy metals of interest

We sought to engineer and characterize transcription factor-based constructs to monitor five heavy metals relevant to marine environments^1,3^: arsenic, mercury, copper, lead, and cadmium. To this end, six constructs were engineered, each consisting of a metal-inducible promoter that drives green fluorescent protein (GFP) expression and, in some cases, its own regulating transcription factor (Figure 1A, Supplementary Table S1). Each construct was designed to produce an output corresponding to the concentration of a specific heavy metal.

**Figure 1.**
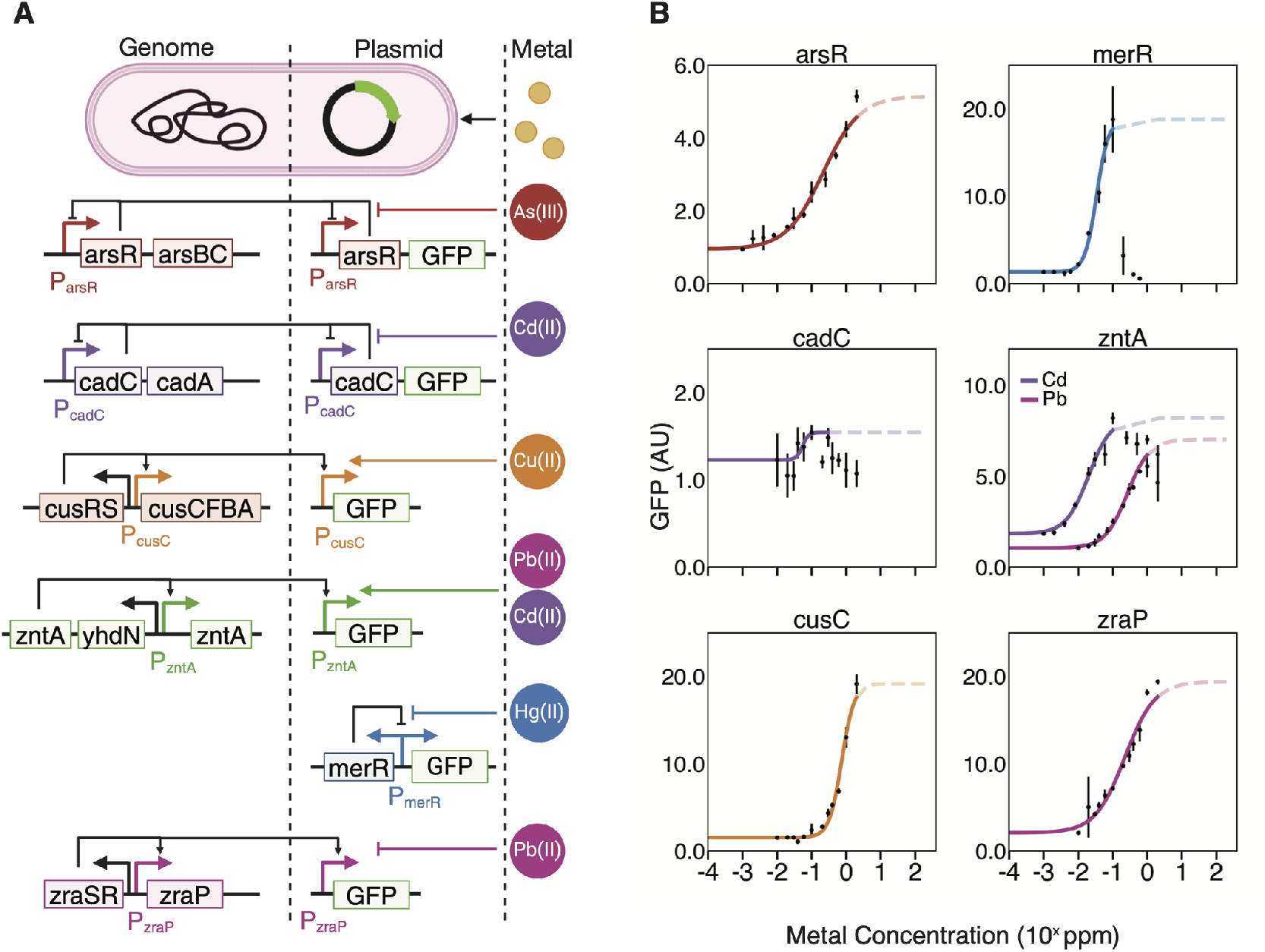
Development and characterization of real-time transcription factor-based biosensors for heavy metal quantification. (A) Six plasmids were engineered to respond to Cu(II), As(III), Pb(II), Cd(II), and Hg(II) (rightmost column) in the *E. coli* K-12 MG1655 strain. The center column indicates the engineered circuit elements that are encoded on the plasmid. In some cases, these engineered elements interact with transcription factors encoded by the host MG1655 genome, as indicated by the leftmost column. Created with BioRender.com. (B) Metalfluorescence dose-response was measured for each of the synthetic fluorescent biosensor strains. Concentrations indicated by the dashed line affect cell growth and were excluded from the dose-response curve fit. Data are *mean ± SD*, with N=3 independent replicates.

All sensor constructs were placed on a medium-copy p15A origin plasmid with kanamycin resistance. The native *E. coli* arsR promoter was selected for arsenic detection. This promoter has previously been shown to respond to arsenic, through the transcriptional repressor protein ArsR, which binds to the promoter region and inhibits transcription in the absence of arsenic^47^. Similarly, the analogous native *E. coli* cadC promoter and transcriptional repressor protein CadC were selected for sensing cadmium^48^. For detection of mercury, we selected the well-characterized MerR transcriptional activator with its corresponding bidirectional promoter^49^. While merR is naturally found in a variety of Gram-negative bacteria, including some species of *E. coli*, it is not naturally found in the MG1655 strain. The promoters cusC, sensitive to copper^50^, and zraP, sensitive to lead^51^, are natively present on the MG1655 genome and were previously found to be responsive without overexpression of additional transcriptional factors. Promoter zntA, responsive to cadmium and lead^52^, was also included, given its well-documented sensor performance.

Sensor construct performance was initially verified in wild-type *E. coli* K-12 MG1655 in minimal media background, with dose-response curves measured using the Hill equation (Figure 1B). Due to the cellular toxicity of heavy metals, several sensors arrest growth before the GFP-toxin response is saturated; however, a detectable level of fluorescence was measured at non-toxic concentrations for all strains. The zntA sensor responded to both cadmium and lead as intended, but its cross-specificity for the two metals revealed a major weakness of single-target WCBs: crosstalk across multiple analytes can lead to confounding signals in mixed samples.

### Multiplexed microfluidics enables efficient dynamic screening and characterization

For further characterization, we developed a versatile, costeffective microfluidic screening system with modular design elements to enable simultaneous testing of multiple microbial strains or multiple conditions. WCBs are typically characterized using single-timepoint approaches, masking dynamic behaviors, which can carry more information on WCB response^9,10,14,18,30,53^. Central to our platform are the cell trapping regions, which are easily interchangeable with various architectures, using our microfabrication process^54,55^. Hydrodynamically trapping cells in a spatially isolated location from the cell deposit spot, the trapping regions feature back channels that allow for even flow of nutrients through the trap^56^ (Figure 2A). Upon device setup, growth media flows through and perpendicular to the spotting region features, carrying spotted cells to the downstream 0.5 µm layer cell traps. Within roughly 8 hours, cell traps reach confluence (Supplementary Figure S1A). The separation of cell spotting and imaging regions generates clear, distinctive dynamics, enabling use of multiple reporting mechanisms (Supplementary Figure S1B). Testing GFP as a reporter, we saw a fast strong readout when induced (Figure 2B). Swapping the reporter with a cell lysis gene, we also demonstrated changes in transmitted light (TL) as another mechanism of dynamic response that is compatible with the hydrodynamic trap geometry (Supplementary Figure S2).

**Figure 2.**
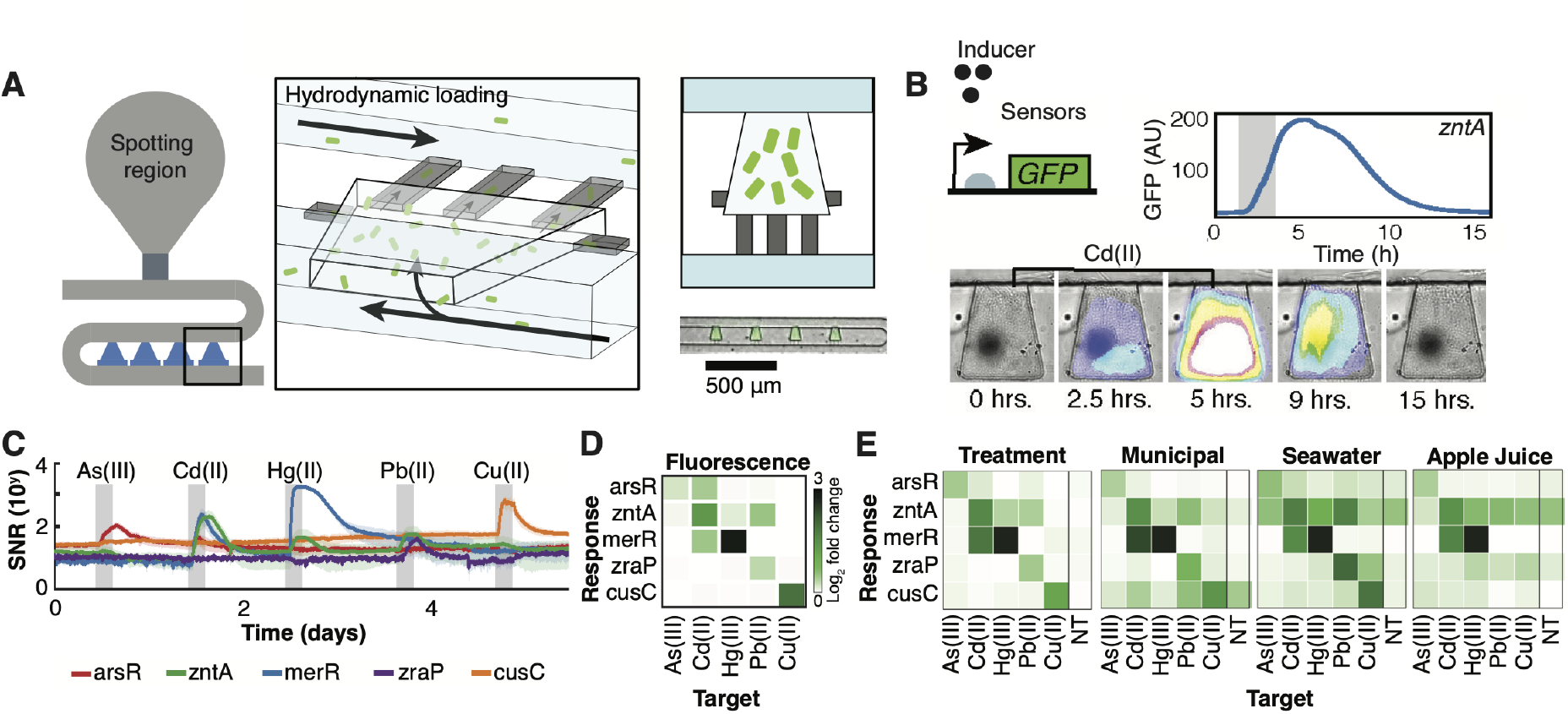
A versatile, multiplexed chemostat array. (A) A drawing of an individual strain’s spotting and trapping region. Each growth region is comprised of a spotting region where cells are deposited, after which hydrodynamic loading enables media flow to carry cells into shallower empty traps for imaging. Upon filling the traps, media flow proceeds through the main channel, perpendicular to the traps. (B) Inducible fluorescence reporting system (GFP expression) tested with our microfluidic layout. The zntA WCB driving GFP is exposed to a two-hour exposure of 1 *µ*M Cd (gray bar). (C) A plot of signal-to-noise ratio of the sensor strains subjected to temporal exposures of the five metals, as represented by the gray shaded regions. Fluorescence traces were extracted over time, and signal-to-noise ratio calculated. Solid lines represent mean of N=12 HD traps in the microfluidic experiment and shaded region represents standard deviation. (D) A heatmap of the fold-change for each fluorescent sensor in response to each heavy metal in the experiment described in panel C. (E) The fluorescent strains were tested on four environmental water sources: water from a local water treatment plant, municipal potable water, seawater, and commercial apple juice. Metal concentrations for panels C-E were: 1 *µ*M As(III), 0.5 *µ*M Cd(II), 0.5 *µ*M Hg(II), 5 *µ*M Pb(II), and 2 *µ*M Cu(II). No toxin (NT) represents an exposure to the water sample with no additional dissolved metals.

To evaluate responses of multiple strains simultaneously, we utilized a scalable manifold array capable of housing 48 strains, based on our previous multiplexed designs^7,56,57^. The 48-strain array maintains a single-layer and single inlet-outlet. Past iterations of this multiplexed platform forewent the hydrodynamic (HD) traps, instead opting for a single trap directly connected to the cell-spotting region^7^. While such an implementation generated useful data, HD traps are spatially isolated from the larger cell mass, smaller, and flanked by media channels on both sides, generating more pronounced dynamics (Supplementary Figure S1B). Furthermore, direct deposition of cells onto our platform circumvents the need for multiple fluidic ports^58–60^and cell loading lines^46,61^typically required to inoculate multi-strain devices without cross-contamination, greatly simplifying the experimental setup of these devices with manual loading capability. This manifold design is easily scalable, allowing for different cell deposition methods (Supplementary Figure S1C-D).

To evaluate a single strain’s response to multiple conditions simultaneously, we transposed our modular HD traps into a single-strain gradient generator device, based on a previously designed branched mixing array upstream of the cell culture region^62^. A discrete gradient of conditions is generated with mixing ratios of two inlet media (Supplementary Figure S3A). This design forewent the cell spotting regions altogether, instead opting for vacuum-loading of cells through the outlet during the final step of the experimental setup. Sulforhodamine dye, an inert dye that does not affect cellular response^63^, was used to visually indicate the gradient across the device as the channels mixed (Supplementary Figure S3B).

The heavy metal WCBs were characterized on our multiplexed 48-strain screening platform and subjected to temporal exposures of each of the five metals (Figure 2C). When exposed to four-hour pulses of each metal, sensors exhibit a typical first order-like response, increasing fluorescence upon metal introduction, with exponential relaxation upon metal removal within 24 hours. The parallel culture of multiple strains on the same device enabled the specific distinction of each metal and the determination of crosstalk during each metal exposure (Figure 2C-D). Interestingly, many of the sensors exhibited off-target responses to heavy metals outside of their intended targets (Figure 2D).

Biosensor response was evaluated on four different water supplies to further assess their real-world deployability. The WCBs were subjected to five consecutive exposures of metals on four different water supplies, spiked with the same heavy metal concentrations as in Figure 2C: water from the Alvarado Water Treatment Plant in San Diego, California which provides drinking water locally, municipal potable tap water, seawater, and commercial apple juice (Figure 2E). On water from the treatment facility, the activation profile for all fluorescent sensors was very similar to the results in lab water. However, on municipal potable water, sensors experienced more crosstalk, especially the cusC and zntA sensors. Apple juice proved to be a difficult sensing background, especially with its high autofluorescence masking the sensor response. The zntA sensor was continuously activated, likely due to the high amount of trace metals, particularly zinc, which is known to interact with the zntR regulator^52^. Furthermore, copper and lead were not detected by zraP and cusC. This may have been attributed to near-toxic levels of trace copper already present in apple juice, the potential presence of chelating agents, or high background fluorescence that obscured the signal. This inconsistency in response in a new background highlighted the weakness of relying on laboratory-characterized single-target sensors. Furthermore, seawater induced an unconventional response in the sensors, likely due to osmolar stress in the high-salinity environment; most laboratory strains of *E. coli* used for biosensor development struggle to survive and function under high salinity environments, suffering hyperosmotic shock and subsequent limitations in population growth^64^. In particular, zntA was activated in the absence of any added metal, despite low levels of trace cadmium and lead in the seawater (Figure 2E). Taken together, the observed inconsistency in sensor responses across various non-laboratory background water supplies revealed that the conventional methods of engineering biosensors for specific analytes under laboratory conditions falter when faced with different and often stressful deployment conditions. Furthermore, disentangling individual signals among combinations of analytes becomes particularly challenging in the presence of crosstalk.

### Adaptive laboratory evolution produces diverse, tractable chassis robust in high salinity environments

We hypothesized that integrating multiple biosensor signals would enhance robustness under deployment conditions where combinations of analytes are present. Although novel chassis using native bacteria are an active area of research to circumnavigate the pressure of withstanding environmental stressors, genetic tractability and circuit behavior portability remain challenging^65–67^. On the other hand, adaptive laboratory evolution (ALE) enables a well-characterized and engineerable strain to efficiently acquire mutations that increase its fitness through prolonged propagations in desired conditions, such as extreme pH or temperatures^68–71^, without *de novo* information for engineering stress tolerance. We aimed to harness the high-throughput, automated capability of ALE to not only enhance the fitness of the *E. coli* K-12 MG1655 laboratory strain in seawater, but also to generate genotypically distinct variants with phenotypic diversity without compromising strain tractability or requiring extensive characterization and tuning.

Previously, MG1655 strains have been evolved to grow rapidly on the non-native substrate sucrose^71^ and in minimal media^70^. As such, we evolved the wild-type MG1655 strain in synthetic seawater to accumulate and select for mutations that improve its fitness under high salinity conditions. Using an automated ALE platform^70,71^, five independent replicates of the MG1655 strain were successively propagated in batch in equal parts minimal media and synthetic seawater (HM9/seawater). The resulting evolved strains, labeled ALE1-ALE5, were isolated from the final populations and whole-genome sequenced (Figure 3A).

**Figure 3.**
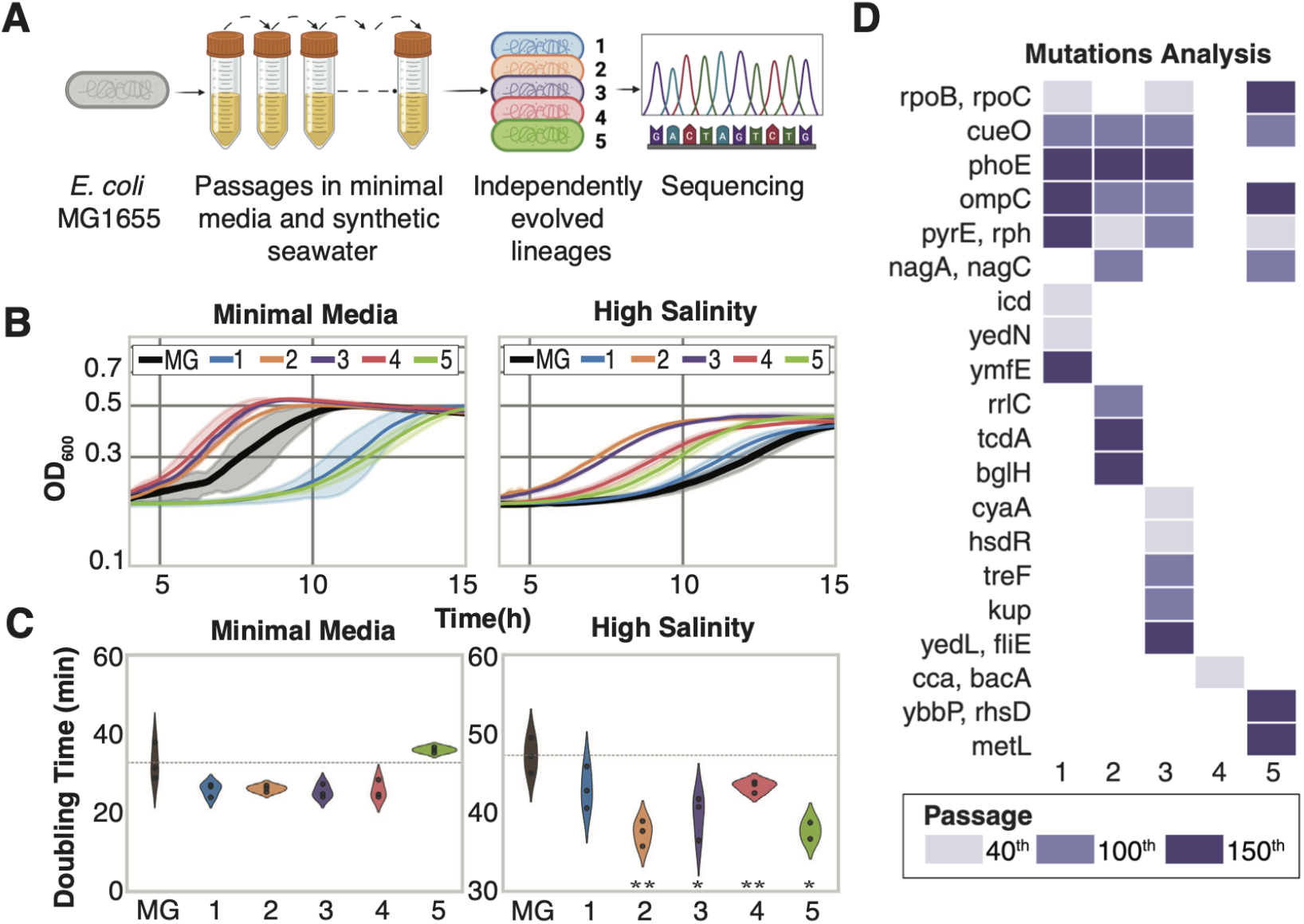
Adaptive laboratory evolution of laboratory *E. coli* K-12 MG1655 to tolerate seawater conditions. (A) The adaptive laboratory evolution process. Five independent replicates of *E. coli* MG1655 were serially propagated in flasks in equal parts minimal media and synthetic seawater (HM9/seawater) using an automated platform. Evolved populations were cryopreserved with every marked increase in growth rate. Final evolved strains were isolated at approximately 150 passages, plated on agar. Created with BioRender.com. (B) Plots of optical density as a proxy for growth for each of the evolved strains in comparison to the starting strain. Solid lines represent mean of N=3 independent wells, and shaded region represents standard deviation. The black line represents the wild-type *E. coli* MG1655 strain. (C) Doubling times in synthetic seawater are reduced for evolved strains, in comparison to the wild-type strain. *P = 0.05 and **P = 0.01. (D) Analysis of genomic changes across different timepoints of ALE process for each strain. Shaded boxes represent presence of mutation in identified gene, and color gradient represents flask number according to the color key.

Evolved strains’ growth rates and doubling times in HM9/seawater were estimated using Curveball, an opensource software used for growth curve analysis from Tecan plate reader measurements^72^ (Figure 3B-C, Supplementary Figure S4A-B). Evolved strains exhibited a statistically significant improvement in doubling time in seawater, with at least 10% reduction below the starting wild-type strain (Figure 3C), suggesting improved fitness to the high salinity environment. This marked improvement in growth of the evolved strains was further validated with NaCl supplementation to minimal media (Supplementary Figure S5). Furthermore, recent studies have indicated lag phase duration as an organized and adaptive process for bacteria, which can vary upon presence of stressors^73^. Under high salinity conditions, the wild-type strain’s lag phase extended almost two hours beyond its evolved counterparts, suggesting its sensitivity to higher osmolarity (Figure 3B).

Sequence analysis of the evolved strains pointed to various mutations that distinguish the evolved strains from the wild-type (Figure 3D, Supplementary Table S2). The pyrE and rph mutations were commonly associated with ALE growth rate selection in minimal media; *E. coli* K-12 wild-types have a frameshift mutation in rph causing pyrimidine starvation, which is typically remedied through the ALE process^70,74^. Apart from these, mutations varied among the evolved strains. Some exhibited phoE and ompC mutations; these are outer membrane porins, which regulate uptake of small molecules^75,76^. To mitigate high salinity stresses, *E. coli* typically employ a strategy of uptake or biosynthesis of biocompatible solutes, such as trehalose or L-glutamate, to counterbalance the flood of potassium ions under high osmolarity conditions^77^. As a result, these mutations could be related to biocompatible solute uptake. Studies in the past decade have suggested that adaptation to stressful environments can occur through loss of function by rewiring the cell’s metabolism without refining enzymatic activities or acquiring gain-of-function mutations^78,79^. As such, observed mutations such as those in nagA and nagC may potentially be knock-outs that confer stress resistance, as suggested in another recent ALE study with osmotic stress^80^. Other mutations between strains were diverse and their effects unclear from sequence analysis alone; however, we postulated that their genotypic diversity gives rise to phenotypic diversity among evolved strains.

### Dynamic screening of biosensors enhances evaluation of diversity in response characteristics

The six engineered plasmids were transferred into the five evolved chassis, producing a set of 30 strains. To evaluate the WCBs’ diversity, they were characterized with our microfluidic screening platforms in HM9/seawater to assess temporal sensor performance diversity. Representative temporal fluorescence data for merR WCBs using the 48-strain array are displayed (Figure 4A-B). Similarly to the wild-type WCBs, four-hour pulses of each metal yielded first-order-like fluorescence responses before relaxing upon metal removal. However, the evolved strains exhibited diverse dynamic response phenotypes, varying in several critical parameters such as maximum fluorescence response, maximum response rate, and rate of relaxation (Figure 4C). The maximum response rate was computed as the maximum of the first derivative of the fluorescence time series. The response relaxation was analogous to voltage relaxation in a resistor-capacitor (RC) circuit charge pulse^81^. As such, the characteristic time constant, ?, was computed as a proxy for the response rate of relaxation. Screening multiple strains simultaneously allowed rapid evaluation of response diversity between evolved strains, making desired biosensor variant selection efficient (Figure 4D). For instance, if particular biosensing needs required rapid response rates, ALE2 may be desirable, given the faster response rates of zntA-ALE2, arsR-ALE2, and cadC-ALE2. Individual temporal data for the other biosensor variants in Figure 4D are available in Supplementary Figure S6.

**Figure 4.**
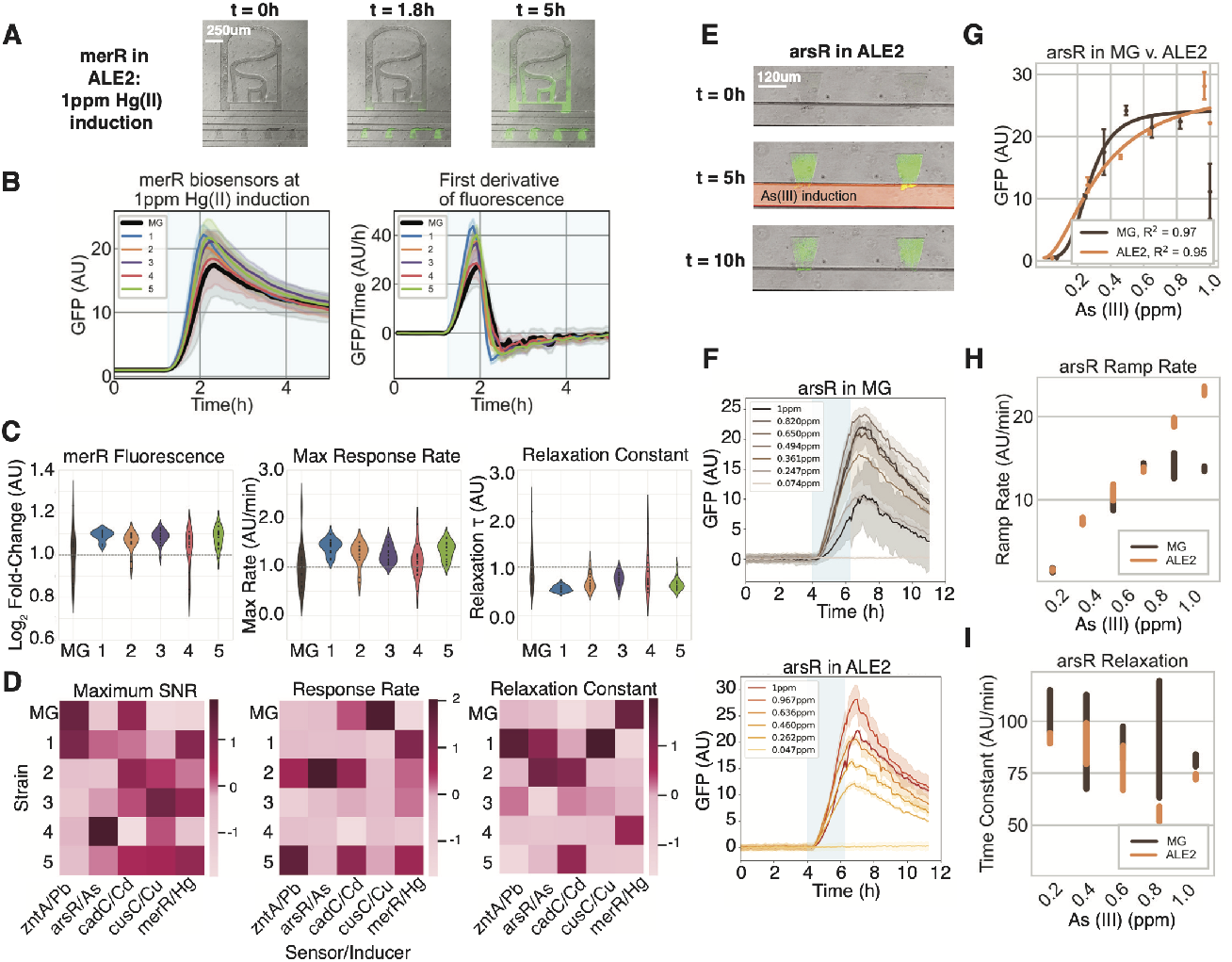
Evolved strains exhibit diverse dynamic behaviors, including fluorescence, response ramp rate, and response relaxation rate in the presence of seawater. (A) A 48-strain multiplexed microchemostat array was employed to simultaneously induce each evolved strain carrying the same heavy metal-sensing plasmid in HM9/seawater. Images of merR-ALE2 induced with 1ppm Hg(II) in multi-strain device are displayed. (B) Fluorescence time traces of each merR biosensor variant, in comparison to merR-MG1655. Solid lines represent mean fluorescence (N=8 HD traps) measured from HD trap regions for each strain, while shading around lines represents standard deviation. Dark line represents the MG1655 WCB, while all other lines represent evolved WCBs. Shading indicates 4-hour induction period. (C) Maximum fluorescence fold-change of each merR biosensor in comparison to merR-MG1655, (left), maximum response rate for each evolved merR biosensor (center), and relaxation time constant for each evolved merR biosensor (right). merR-MG1655 is normalized to 1 for all metrics. (D) Maximum signal-to-noise ratio, maximum response ramp rate, and relaxation time constant for each biosensor strain in the presence of the indicated heavy metal. Respective units of measure are the same as in (C). (E) Images from microfluidic experiments before, during, and after induction of ALE2 carrying arsR. Sulforhodamine dye (red) was used as a visual indicator for the presence of heavy metal upon induction. (F) MG1655 and ALE2 carrying arsR biosensor, exposed to simultaneous gradient of As(III) concentrations. The blue shaded box indicates when heavy metal is introduced. Solid lines represent mean fluorescence (N=4 HD traps) measured from HD trap regions for each strain, while shading around lines represents standard deviation. (G) Hill-curve fit of MG1655 and ALE2 carrying arsR plasmid. (H) Maximum response ramp rate of MG1655 and ALE2 arsR strains at each As(III) concentration. (I) Relaxation time constant of MG1655 and ALE2 arsR strains at each As(III) concentration.

We further demonstrated our modular microfluidic platform by using our single-strain gradient generator to efficiently screen biosensor responses to multiple conditions in parallel. This device was employed to examine the responses of the arsR-MG1655 and arsR-ALE2 biosensors to multiple arsenic concentrations simultaneously (Figure 4E,Supplementary Movie 1). Sulforhodamine dye was used to visually indicate the 1 ppm arseniccontaining HM9/seawater, as the channels mixed it with plain HM9/seawater media. The temporal GFP responses of the two strains are displayed for each concentration generated by the device (Figure 4F). Using the maximum fold-change responses from the temporal data, a Hill function was fitted Figure 4G). The Hill curves revealed that fluorescence response monotonically increased in correspondence with arsenic concentrations for a wider concentration range in arsR-ALE2 than in the wild-type strain, with arsR-MG1655 reaching its maximum at approximately 0.494 ppm and arsR-ALE2 at 0.967 ppm, nearly double. The concentrations where a drop-off in fluorescence occurred were omitted from the Hill curve fit, as the responses were no longer reflective of increasing metal concentration. By using the single-strain gradient device, this difference in monotonic response range was rapidly illuminated.

Beyond GFP fold-change, other dynamic characteristics, including maximum response rate and relaxation rate, were observed, similarly to the multi-strain device (Figure 4H-I). For relaxation rate, concentrations yielding no significant response (defined as below 1 AU) were excluded. These characteristics varied with induction concentrations. For example, the response ramp rate increased monotonically for the wild-type strain up to approximately 0.6 ppm, despite its fold-change capping at approximately 0.5 ppm; therefore, the response rate provided an additional metric beyond single-timepoint measurements. Taken together, the multiplexed characterizations of the WCBs with high temporal resolution demonstrated the phenotypic diversity across the evolved strains, achieved through ALE with no additional rational engineering. This response diversity could provide more information in the presence of a combination of analytes in deployment conditions than single-target laboratory-characterized WCBs.

### Rich response diversity across evolved strains facilitates combinatorial prediction

A longstanding challenge of WCBs is their difficulty with combinatorial detection, as most samples contain multiple targets, causing biosensor crosstalk and complicating accurate detection^42^. As such, a biosensor system that could sense a combination of analytes in parallel would be economical and efficient. Supervised learning classification approaches have recently expanded biological data processing capabilities^82–85^. Using the observed rich diversity of biosensor response phenotypes, we constructed a consortium biosensor and applied a supervised learning classifier, benchmarking its predictive accuracy against that of the single-target WCBs in classifying a combination of all five heavy metals simultaneously.

As a proof of concept, we used batch culture plate reader experiments to characterize features of the evolved strains. Though these did not provide dynamic features, they were insightful in demonstrating combinatorial capabilities in high-throughput. As such, the evolved WCBs were exposed to different concentrations of each heavy metal to evaluate their dose-dependent responses in HM9/seawater (Figure 5A, Supplementary Figure S7). The diverse response profiles offered a selection of sensors that can be chosen depending on the desired biosensing applications.

**Figure 5.**
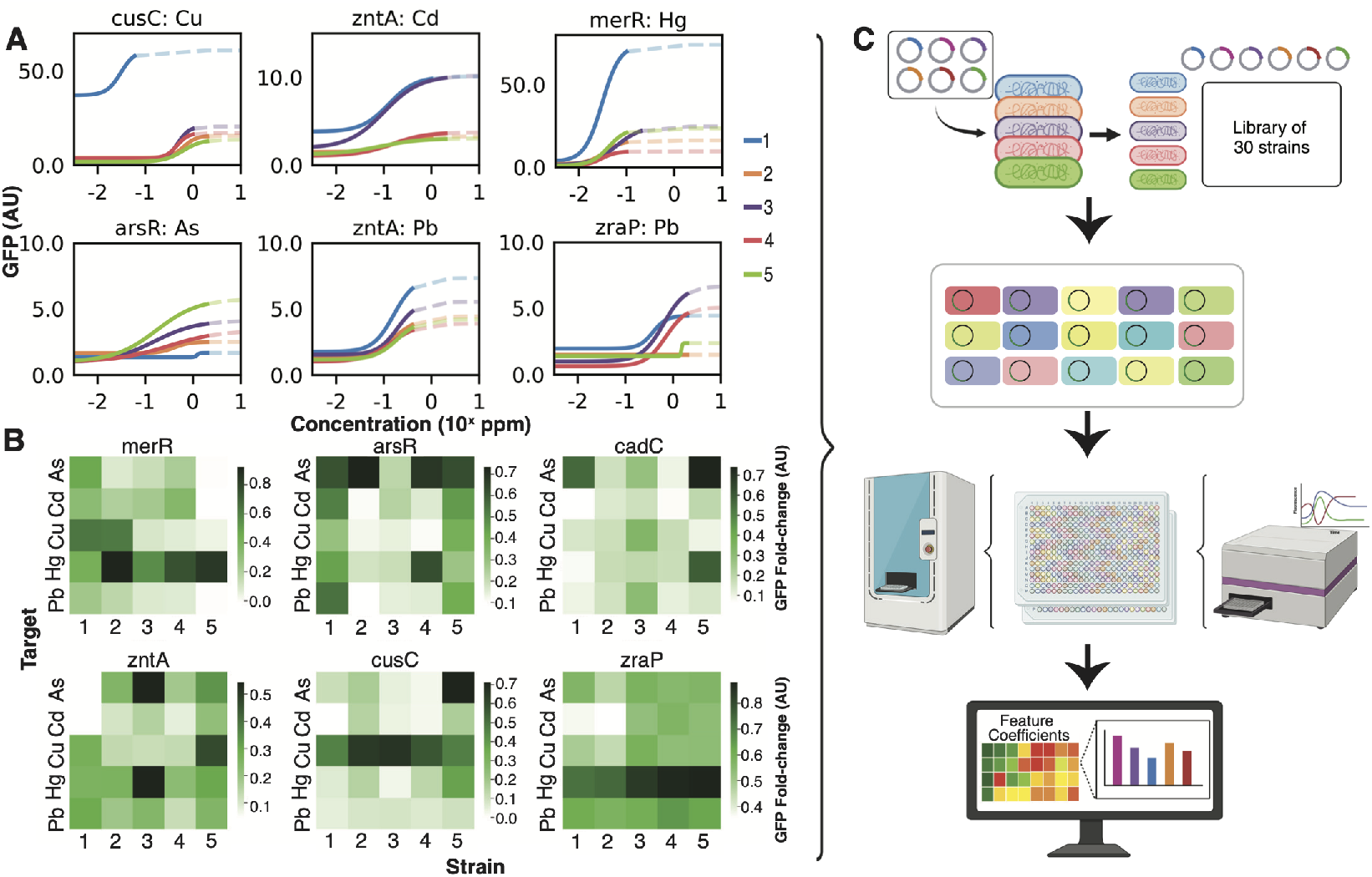
Response diversity across evolved strains. (A) Dose-response curves of each evolved biosensor strain in HM9/seawater when exposed to the heavy metal it was engineered to detect, showcasing the strains’ diversity. Data shown are mean, with N=3 independent replicates, and solid and dashed lines are as indicated in Figure 1B. (B) Characterization of specificity across evolved strains indicated potential for machine learning classification due to their diversity. Each strain, MG1655 and evolved versions, carrying each sensor plasmid, was exposed to each heavy metal. The color gradient corresponds to each sensor variant’s fold-change response to the corresponding metal. Heavy metal induction concentrations are the same as in Figure 2D-E. (C) Schematic illustrating development of a training set for a supervised learning classification model. A collection of 15 strains was selected and exposed to approximately 500 heavy metal combinations, resulting in a rich training set for developing a machine-learning model. Created with BioRender.com

Furthermore, we examined the evolved WCBs’ fold-change responses to each heavy metal to evaluate cross-specificity in HM9/seawater (Figure 5B). Interestingly, the evolved strains, even when carrying the same plasmids, did not display the same levels of specificity or cross-specific interactions. In fact, some of the WCBs exhibited greater responses to heavy metals they were not designed for. For example, merR-ALE1 and arsR-ALE1 became responsive to the other metals. Most of the zntA biosensors displayed promiscuity, similarly to the zntA-MG1655 biosensor, likely because zntA is a P-type ATPase that works as a pump for multiple divalent heavy metals, including lead, cadmium, and zinc^86,87^. Notably, the cadC biosensors exhibited a response to cadmium with weak specificity, but some of strains cadC-ALE5, responded strongly to other metals, such as mercury and arsenic. The zraP biosensors showed strong specificity to mercury and high promiscuity.

Instead of treating multi-target specificity as a hindrance as in conventional single-target biosensors^42,88^, we employed a strategy to harness the combined responses of multiple strains to combat crosstalk effects and produce more robust predictions. By examining the combined phenotypic behaviors of selected biosensors in parallel, we could generate a distinctive response signature, similar to a barcode or a fingerprint, to quantify individual metals in a combined sample. As such, a more compact set of 15 WCBs was selected from the full set to test combinatorial predictive capability (Figure 5C). Three variants for each heavy metal with varied specificity and sensitivity were selected. Strains cadC-ALE1, cadC-ALE3, and cadC-ALE5 were included, since they exhibited varied but clear specificity to heavy metals outside of cadmium, though cadmium was their intended target. The zraP WCBs were excluded due to their strong responses to mercury and high promiscuity, since most of the merR WCBs already exhibited strong specificity to mercury.

Using an ECHO acoustic droplet handler, approximately 500 heavy metal combinations were introduced to each of the 15 biosensors to collect a training dataset (Figure 5C). The combinations included different concentrations of each of the five heavy metals. Features were extracted for proof-of-concept, including fluorescent fold-change response maxima and minima, number of peaks, and curve skewness, to generate a response signature (Supplementary Figure S8A). Supervised machine learning classification models were trained on these response signatures from the 15 WCBs in response to the heavy metal combinations to quantify the concentrations of the heavy metals present^89^. Multiple supervised learning classification models were applied and compared, including polynomial regressions, decision trees, and ordinary least squares regression, using the scikit-learn Python packages. A standard loss function was used as a comparison metric of the predictive strengths of each classifier (Supplementary Figure S8B). Using a k-fold cross-validation strategy to prevent model bias and overfitting, the training set of combinations was divided into k=6 segments, each of which served as a validation set while the model was trained on the other five segments. This process was repeated, with each of the six segments serving as the validation set. The loss function across the k-folds was averaged for each model. The ordinary least squares model proved sufficient. One major disadvantage and critique of complex machine learning models is that they tend to be a “black box”, masking how the model makes decisions for the user^90^. However, the ordinary least squares model provides information by revealing the magnitude of the feature coefficients, elucidating its pathway to identifying each heavy metal concentration (Supplementary Figure S8C).

To evaluate the predictive capabilities of the model, the consortium biosensor’s performance was benchmarked against the single-target biosensor paradigm, using the same loss function. A test dataset was collected with more than 100 new unseen heavy metal combinations, and the consortium model built using the training set was applied on this new test set. To evaluate the single-target biosensor predictions, each individual WCB’s predictions for its intended heavy metal target was computed using its corresponding dose-response data (Supplementary Figure S9). From these predictions, the most optimal single-target WCB for each of the five heavy metals was identified and compared to the consortium-based predictions (Figure 6A). The consortium-based predictions outperformed the single strain predictions of the most optimal individual sensors in 93% of the sample combinations, exhibiting significantly reduced total loss in using the consortium’s responses (Figure 6B-C, Supplementary Figure S10). We further analyzed the consortium-based predictive performance against those of the single-target sensors for each of the heavy metal constituents (Supplementary Figure S11). For most of the heavy metals, with the exception of cadmium, the consortium outperformed the predictions of the single-target sensors in a majority of the sample combinations: 68% for mercury, 78% for lead, 90% for arsenic, 46% for cadmium, and 86% for copper. Finally, the proportion of false negatives generated by the consortium-based and single-target sensors for each heavy metal was computed, as false negatives can be deleterious when identifying whether a heavy metal is present at all. For most of the heavy metals, outside of mercury, the consortium-based sensor exhibited reduced false negative rates (Supplementary Figure S12).

**Figure 6.**
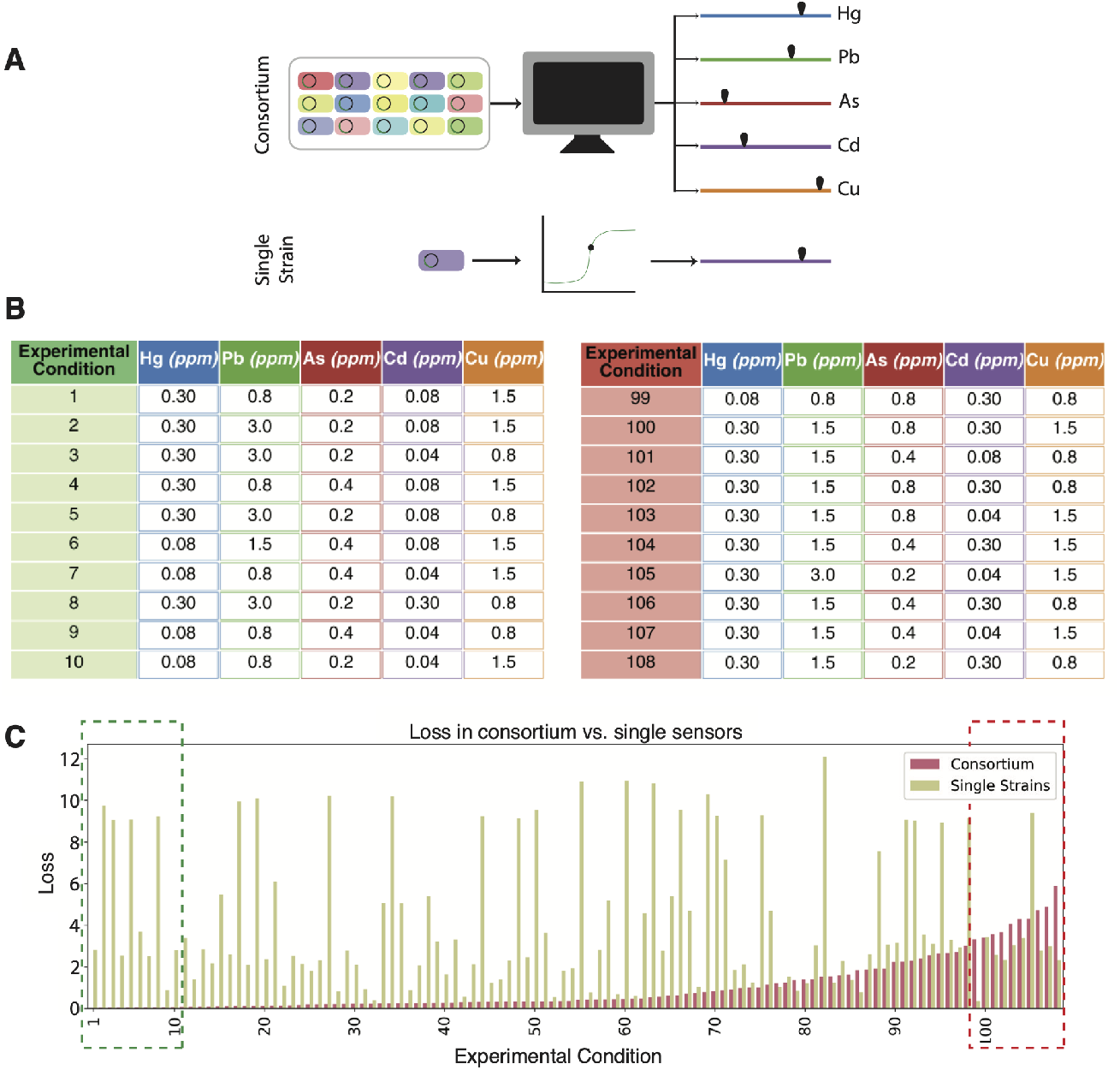
Consortium outperforms single-target biosensors in predictions of heavy metal combinations. (A) Schematic illustrating response from consortium vs. conventional single strain responses.(B) Sample experimental conditions, including the conditions in which the model performed the best and the worst. (C) Comparison of total loss across all heavy metal predictions according to loss function between consortium and single-target biosensor predictions, organized in order of increasing loss in consortium predictions. Heavy metal combinations of the consortium’s best and worst predictions are specified in the tables.

To further evaluate the consortium predictions, we separated the total loss into its individual heavy metal constituents to analyze each metal’s loss contributions (Supplementary Figure S13A). This deconvolution revealed that mispredictions in lead concentrations consistently contributed a majority of the loss in the conditions where the model faltered and produced greater loss. This loss is further exemplified in the absolute differences between the predicted and true concentrations across the experimental conditions (Supplementary Figure S13B-C). With the exception of lead, the consortium predictions were mostly within 0.5 ppm of the true concentrations. Mercury and cadmium were the most closely predicted, with arsenic and copper consistently slightly underestimated. Lead predictions were vastly varied, which is likely due to the promiscuity of zntA, the primary lead sensor in the consortium. Further improvement of the consortium model could potentially be achieved by inclusion of more WCBs with diverse responses to lead, such as the excluded zraP biosensors, or tuning the lead sensor plasmid construct itself to produce more specific responses.

## Discussion

As whole-cell bacterial biosensor development has focused on detecting targets in isolation under laboratory conditions, we engineered a consortium biosensor to combinatorially quantify multiple heavy metals in seawater. We employed adaptive laboratory evolution to obtain a diverse range of *E. coli* strains optimized for high-salinity seawater environments, while bypassing the need for additional rational engineering. Housing transcription factor-based biosensing constructs for the detection of five heavy metals of interest in these variants, the final WCBs were screened using a multiplexed, modular microfluidic platform. This measurement of dynamic temporal behavior highlighted the diversity achieved through ALE. Using the diversity in sensor behaviors and cross-interference collectively as a “fingerprint” readout, we applied supervised machine learning to quantify combinations of heavy metals in seawater conditions. Toward environmental monitoring, the tools developed in this work tackled some of the pressing issues in biosensing, including design for stressful environments, capture of dynamic output, and disentanglement of multiple signals amidst crosstalk.

WCBs have previously been developed for the detection of harmful compounds for environmental safety^58,91–97^, metabolites relevant to bioproduction^98–102^, or biomarkers in *in vivo* medical diagnostics^103–106^. However, engineering for deployability requires sensors robust to the harsh environments of their application^17^. Adaptive laboratory evolution provides a rapid method for achieving resilient variants^68–71^. Future work on WCBs could utilize ALE to generate strains for extreme temperatures, pH differences, or hypoxic conditions. Recent advances in bioinformatics has also opened the door to probing the genomic mutations for characteristics essential to fitness in specific environments, informing future rational engineering approaches^68–70^.

Furthermore, well-established analytical techniques such as atomic absorption spectroscopy (AAS) or inductively coupled plasma mass spectrometry (ICP-MS) are highly sensitive but are limited not only by the number of detectable compounds^95–97^, but also by measurement of a single sample at a single timepoint^107,108^. Multiplexed microfluidic platforms enable screening of multiple engineered strains to multiple conditions while collecting their dynamic data in real time^58,61^. We believe the accessibility, throughput, dynamics, and week-long timescales achievable with our microfluidics screening tool can push the frontier of the biosensing landscape. More broadly, the microfluidics platform can serve as a useful tool to screen genetic circuit outputs in parallel for a desired phenotype, closing the gap between our rapid generation of genotypic diversity and our ability to screen complex phenotypes^109^. Microfluidics has served as a powerful tool for approximating complex real-world environments, spanning soil to human organs^110,111^. While not a perfect recreation of these complex environments, tuning environmental and time-dependent parameters achievable with microfluidics serves an important role in the prototyping and scale-up of all classes of gene circuits. Despite limited means for dynamic phenotype screening, canonical gene circuit motifs, including oscillators, logic gates, and feedback controllers, have been increasingly deployed in time-dependent applications spanning metabolic engineering to therapeutic delivery^20,112–115^. Multiplexed microfluidics, such as ours, can aid in the development of circuits like these.

Typically a hindrance rather than an asset^42^, the crosstalk generated by the diverse strains was incorporated into a supervised learning classification model to quantify multiple heavy metal inputs. Outperforming the single-target biosensor predictions, the consortium biosensor’s promising performance advances the ability to combinatorially detect multiple targets in real-world samples. Harvesting the feature coefficient information could help identify features and strains most useful to the model, helping to narrow down a minimal set of strains or identify the most essential features for a particular biosensing application. Future approaches that could further enrich the field of biosensing include integrating dynamic features, such as frequency measurements, response rate, and relaxation rate, such as those collected in our microfluidics platform. Particularly when combined with newer, more advanced machine learning classifiers, such as neural networks, the consortium biosensor paradigm can enable even more refined predictions.

## Acknowledgments

We thank S. Kumar for taking the time to read the manuscript and providing constructive critiques and feedback. A.J.C. was supported in part by the National Science Foundation Graduate Research Fellowship Program under grant number DGE-2038238. N.H. and J.H. were supported by R01GM144595 and R01AG086348. We acknowledge the Novo Nordisk Foundation (NNF20CC0035580) for funding for the project.This work was supported by the U.S. Department of Energy, Office of Science, Office of Biological and Environmental Research, Genomic Science Program under Secure Biosystems Design Science Focus Area IMAGINE BioSecurity: Integrative Modeling and Genome-scale Engineering for Biosystems Security under contract no. DE-AC36-08GO28308.

## Author contributions

A.J.C. and J.H. designed the project. A.J.C., M.F., N.H., K.Z, and J.H. wrote the paper. A.M.F. designed the adaptive laboratory evolution experiments and interpreted the results. E.C.O. and R.M. executed the adaptive laboratory evolution experiments. N.C., R.O., and E.S. designed and tested the multi-strain microfluidic screening devices. A.J.C. and A.D. designed the single-strain gradient microfluidic device. L.B. designed the biosensor plasmids. P.N. provided guidance for implementing the supervised machine learning classification. A.J.C. and M.A. performed and analyzed the evolved biosensor and high-throughput experiments.

## Methods

### Method details

#### Biosensor plasmid construction

Biosensing plasmids were constructed using PCR amplification of host strain components listed in Supplementary Table S1, followed by Gibson Assembly. The green fluorescent protein-containing backbones were obtained from existing plasmids in our lab. Plasmid sequences were confirmed with Sanger sequencing (Integrated DNA Technologies, San Diego, CA) and transformed into chemically competent *E. coli* K-12 MG1655 cells and evolved variants.

#### Plate reader experiments

Overnight cultures grown in LB media were reseeded into HM9 minimal media or 1:1 HM9 to synthetic seawater (HM9/seawater) at 1:1000 dilution and grown until they reached an OD_600_ of about 0.3-0.4. The HM9 minimal media composition, containing minimal traces of metal, was based on prior studies^7,116^ and optimized for microfluidic *E. coli* growth. For fluorescence dose-response curves, 96-well plates were prepared with media and varying heavy metal concentrations. Cultures were diluted to OD_600_ of about 0.1 on the plate. Fluorescence and OD_600_ values were measured using an Infinite® M200 PRO Multimode Microplate Reader. For high-throughput experiments, 384-well plates were used, and inducer was transferred to the plates using the Echo Acoustic Liquid Handler. For all plate reader experiments, fluorescence measurements were taken at maximum induction, which occurred around 2 hours into the experiment. This was normalized by fluorescence of a promotorless strain to correct for cell autofluorescence and by OD_600_.

#### Microfluidic device development and fabrication

Microfabrication techniques in patterning SU-8 photoresist onto a silicon wafer to create a mold for microfluidic devices have previously been described^54,55,57^. Polydimethylsiloxane (PDMS) microfluidic devices were made with a mixture of 10:1 Sylgard 184 to curing agent, which was poured over the wafer centered on a glass petri dish and degassed to remove bubbles in the PDMS. The wafer with the PDMS was cured for one hour at 95°C on a level surface.

#### Microfluidic device loading and bonding

PDMS devices and glass slides were cleaned with MilliQ and 70% Ethanol. Adhesive tape was used to remove any remaining fibers on the device. Overnight cultures were reseeded at 1:100 to an OD_600_ of approximately 0.3-0.4. For multi-strain devices, 1mL of the overnight cultures was centrifuged, the supernatant was discarded, and cell pellets were resuspended in 20 µL fresh LB to make highly concentrated cell cultures. The devices and slides were exposed to oxygen plasma, after which 23 ga syringe tips were used to transfer cells from the concentrated suspension to the devices by lightly contacting the cell traps on the PDMS. The device and glass slide were then bonded and cured at 37°C for two hours. To load the gradient devices, they were instead bonded and cured prior to vacuum loading using a previously described protocol^62^.

#### Microfluidic experimental protocol

Microfludic experiments were performed on a Nikon TE2000-U epifluorescent inverted microscope (Nikon Instruments Inc., Tokyo, Japan). The inlet and outlet ports were connected to 50mL syringes and Tygon tubing. The inlet syringes were filled with HM9/seawater with Kanamycin and 0.075% Tween-20. Tween-20 is a surfactant to help prevent biofilm formation that can clog microfluidic channels and reduce the longevity of experiments. This has previously been shown to have no adverse effect on *E. coli* in many microfluidic experiments^55,92^. The waste outlet was filled with approximately 5 mL of MilliQ water, to provide negative pressure. The inlet and outlet syringes were separated by a 10” height difference. Strains were grown to confluence in the cell traps after at least 8 hours, and a predetermined heavy metal concentration was pipetted into the inlet syringe. After a 4-hour induction period, any remaining inducer solution was pipetted out and replaced with fresh HM9/seawater. Fluorescence time series were extracted and normalized to remove device background fluorescence and strain background fluorescence.

#### Source water testing

Sensor strains were tested on four different water sources: (1) water from the Alvarado Water Treatment Plant in San Diego, CA, (2) municipal potable tap water from San Diego, CA, (3) Seawater from Blacks Beach in La Jolla, CA (3) and apple juice (Trader Joe’s Fresh Pressed Apple Juice SKU#88463). Once a day, each device was exposed to a mixture of a single toxin dissolved in HM9 made from the respective water source for four hours. Signal-to-noise ratio was determined as the fluorescence value at each time point divided by the standard deviation of fluorescence 7.5 hours before induction. Fold change was calculated from the signal-to-noise ratio for four biological replicates of each sensor.

#### Adaptive laboratory evolution

Adaptive laboratory evolution (ALE) was carried out by Adam Feist using a previously described method^70,71^. Briefly, five biological replicates from *E. coli* strain K-12 MG1655 were grown overnight in HM9/seawater. Using an automated liquid-handler, this culture was serially passaged during exponential phase over approximately 140 passages into fresh HM9/seawater. Final strains used in experiments were isolated at the end of the evolution process and whole-genome sequenced.

#### Batch growth assays

Overnight cultures were grown in LB media at 37°C with antibiotics for 16 hours and reseeded into 1:100 diluted HM9/seawater media. Once an OD_600_ of approximately 0.3-0.4 was reached as measured by the Spectrophotometer, they were diluted to OD600 of approximately 0.01. Growth was then monitored by measuring OD_600_ were taken every 5 minutes for 12-16 hours.

#### Supervised learning classification

The scikit-learn machine learning package in Python was used for training and testing the different classifier models, including polynomial regression, ordinary least squares, and decision trees. GFP fluorescence of each strain was normalized by GFP fluorescence with no inducer added. Features were extracted from each time trace, including the maximum fluorescence, minimum fluorescence, time at which the fluorescence reaches half its maximum, number of peaks, and skewness of curve. These features were standardized to prevent feature bias.

#### Quantification and statistical analysis

To calculate the normalized fluorescence of multi-strain microfluidic experiments, the mean fluorescence pixel value (GFP) for each trap was extracted across the 10X microscope image stack using ImageJ. All GFP values were background-normalized to the mean GFP pixel value in a selection of equal size in a part of the chip with no cells (GFP_*BG*_). Background-normalized GFP values were normalized to the inverse of similarly background-normalized transmitted light 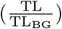 pixel values, as a proxy for cell density. The following formula was used:

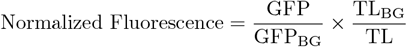

