## Supplemental Information for "Evolved microbial diversity enables combinatoric biosensing in complex environments"

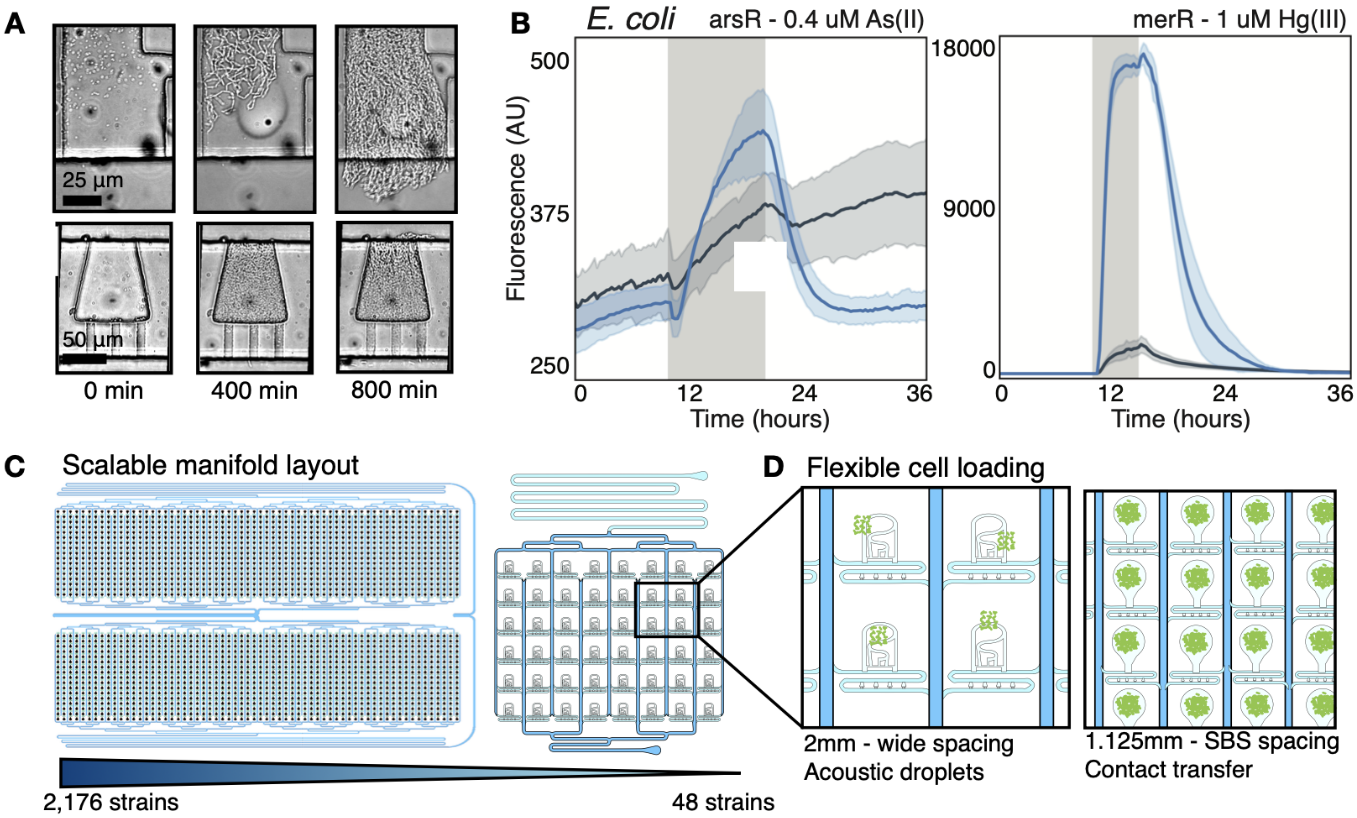


Figure S1: **Microfluidic device scalability and flexibility for cell transfer.** (A) HD traps fill within 8 hours of loading. (B) Direct comparison of fluorescence traces extracted from the spotting region (dark gray) and HD traps (blue). Each strain was grown in the absence of the inducing metal, and exposed for a several hour induction window (gray bars). (C) Using branched, parallel fluidic manifolds, multiplexed arrays can be scaled to house between 48 and 2,176 individual strains [7, 117]. (D) Numerous cell loading methods are compatible with the manifold design, including acoustic droplet ejection (2 mm between strains) of liquid cultures and contact transfer using pin pads of colonies from solid agar plates (SBS-spaced, 1.125 mm between strains). Spotting region geometry and spacing can be adjusted to best fit the spotting method.

| Promoter | Transcription Factor | Source Strain |
| --- | --- | --- |
| $P_{arsR}$ | arsR | *E. coli* R773 |
| $P_{cadC}$ | cadC | *S. Aureus* pI258 |
| $P_{merR}$ | merR | Tn21 |
| $P_{cusC}$ | n/a | *E. coli* MG1655 |
| $P_{zntA}$ | n/a | *E. coli* MG1655 |
| $P_{zraP}$ | n/a | *E. coli* MG1655 |

Table S1: **Biosensor plasmid construct parts.** Each biosensor plasmid is constructed of a heavy metal-responsive promoter, which is regulated by a transcription factor. In $P_{arsR}$, $P_{cadC}$, and $P_{merR}$, the regulatory transcription factors are included in the plasmid, because it is borrowed from a different host strain. In $P_{cusC}$, $P_{zntA}$, and $P_{zraP}$, the regulator transcription factors the plasmid interacts are endogenously expressed in the *E. coli* K-12 MG1655 host strain.

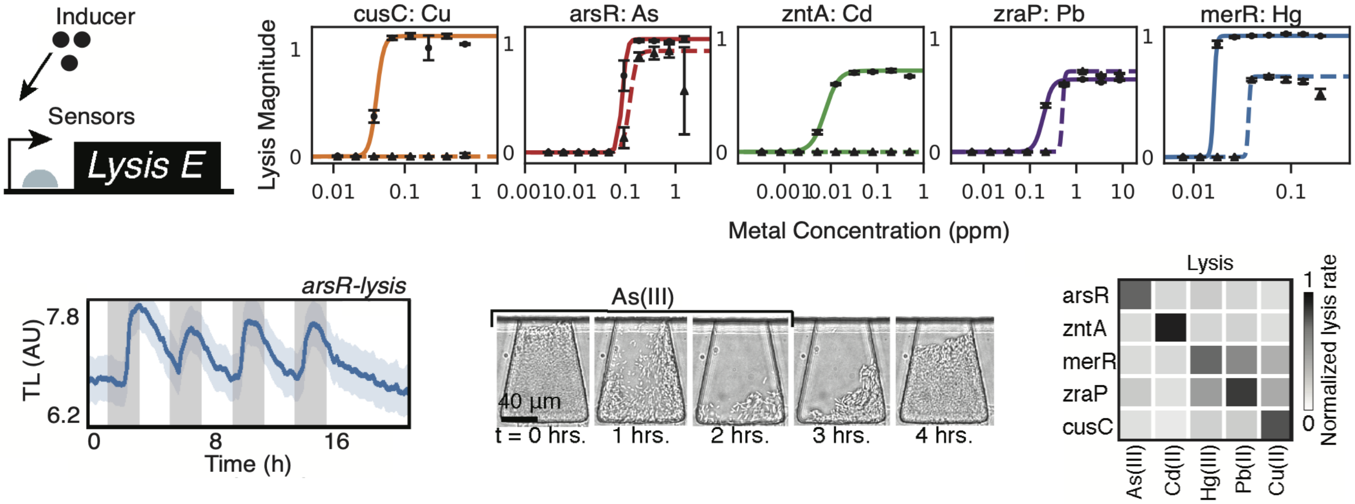


Figure S2: **Biosensor constructs with lysis reporting.** (A) A lysis-based inducible reporting system using transmitted light changes (E lysis protein from phage $\phi$X174). (B) Lysis magnitude is measured as the ratio of the change in OD due to lysis over the change in OD due to growth before toxin exposure. Lysis magnitude is measured at 1 hour (solid line, black circles) and 8 hours after toxin exposure (dashed line, black triangles). Error-bars represent standard deviation of triplicate samples. (C) Lysis-based circuit responding to periodic 2-hour pulses of 75 ppb As(III) (gray bars). (D) Fold change of each lysis sensor when run on microfluidics are shown in response to each heavy metal.

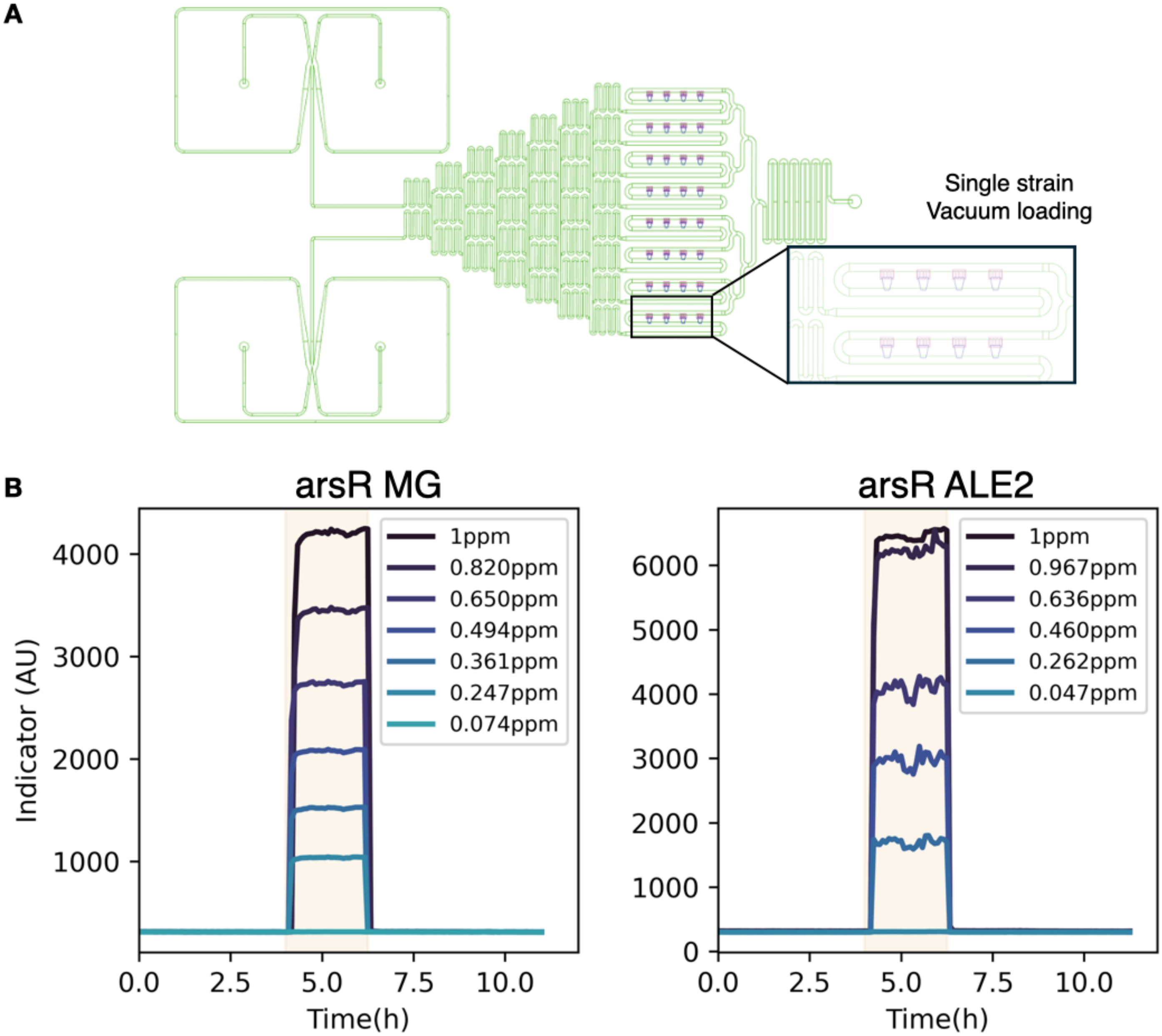


Figure S3: **Gradient generator with HD traps.** (A) Microfluidic device containing HD traps configured to generate an inducer gradient for single-strain characterization across multiple conditions in parallel, based on a previous gradient design [62]. Cells can be rapidly vacuum-loaded and the gradient is automatically generated using the branched channel network. (B) Gradient in concentrations as measured by sulfurhodamine dye, simultaneously generated by the microfluidic device. Shaded area indicates the period of induction, and color gradient from light to dark mirrors the increase in concentration of As(III). Solid lines represent mean fluorescence (N=4 HD traps) measured from HD trap regions for each strain.

**A**

$$\frac{dN}{dt} = rN\left(1 - \frac{N}{K}\right)$$

$$N(t) = \frac{K}{1 - \left(1 - \frac{K}{N_0}\right)e^{-rt}}$$

N = population size
$N_0$ = initial population size
r = initial per capita growth rate
K = maximum population size

**B**

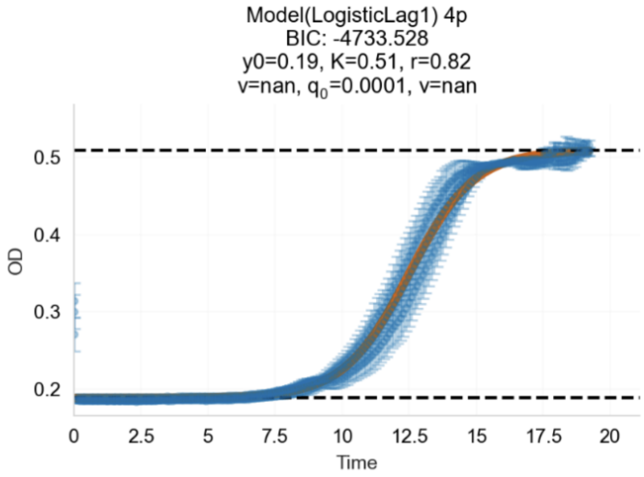


Figure S4: **Growth rate calculation using Curveball.** (A) Logistic model Curveball uses to estimate the growth rate. (B) Example output from Curveball logistic model.

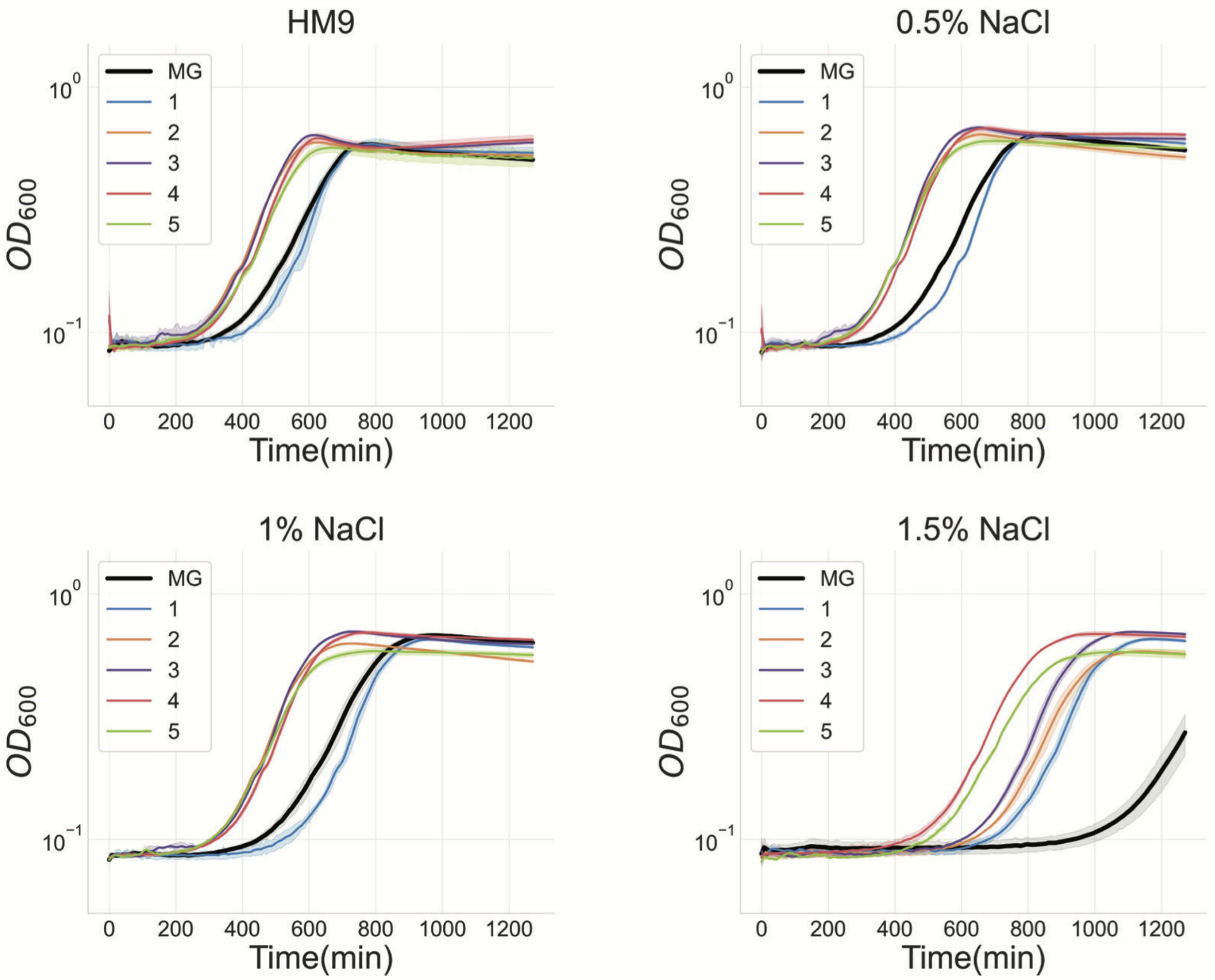


Figure S5: **Growth curve controls with added NaCl.** Growth curves of each evolved strain at different concentrations of NaCl. Solid lines represent mean of N=3 independent wells, and shaded region represents standard deviation. The black line represents the wild-type *E. coli* MG1655 strain.

| Position | Mutation Type | Sequence Change | Gene (Scrollable) | Details |
|---|---|---|---|---|
| Strain 1 | | | | |
| 138,616 | MOB | IS5 (+) +4 bp | cueO | coding (1534-1537/1551 nt) |
| 1,196,220 | SNP | C->T | icd | H366H (CAC->CAT) |
| 1,196,232 | SNP | C->T | icd | T370T (ACC->ACT) |
| 1,196,245 | SNP | T->C | icd | L375M (TTA->CTG) |
| 1,196,247 | SNP | A->G | icd | L375M (TTA->CTG) |
| 1,196,277 | SNP | C->T | icd | N385N (AAC->AAT) |
| 1,196,280 | SNP | G->C | icd | A386A (GCG->GCC) |
| 1,196,283 | SNP | A->G | icd | K387K (AAA->AAG) |
| 1,196,292 | SNP | C->T | icd | T390T (ACC->ACT) |
| 1,196,301 | DEL | Δ15,100 bp | icd,ymfD,ymfE,lit,intE,xisE,ymfH,ymfI,ymfJ,ymfK,ymfT,ymfL,ymfM,oweE,ymfN,aaaE,ymfR,beeE,jayE,ymfQ,ycfK,tfaP,tfaE,stfE,pinE,mcrA | |
| 2,312,011 | INS | (+) T | ompC | coding (739/1104 nt) |
| 3,815,810 | DEL | Δ1 bp | pyrE, rph | intergenic (-42/+53) |
| 4,182,809 | SNP | C->A | rpoB | S522Y (TCT->TAT) |
| 4,183,814 | SNP | T->A | rpoB | V857E (GTG->GAG) |
| Strain 2 | | | | |
| 138,575 | SNP | C->A | cueO | A498E (GCG->GAG) |
| 260,264 | MOB | IS5 (-) +4 bp | phoE, proB | intergenic (-164/-121) |
| 702,331 | DEL | Δ1 bp | nagA | coding (421/1149 nt) |
| 2,312,333 | MOB | IS2 (-) +5 bp | ompC | coding (413-417/1104 nt) |
| 3,815,810 | SNP | G->T | pyrE, rph | intergenic (-42/+53) |
| 3,945,587 | SNP | G->T | rrlC | noncoding (1884/2904 nt) |
| Strain 3 | | | | |
| 138,611 | MOB | Δ3 bp :: IS186 (-) +7 bp | cueO | coding (1529-1535/1551 nt) |
| 2,312,239 | SNP | G->A | ompC | Q171* (CAG->TAG) |
| 3,440,459 | SNP | G->A | rpoA | R191C (CGT->TGT) |
| 4,186,605 | SNP | A->C | rpoC | H419P (CAC->CCC) |
| Strain 4 | | | | |
| 137,918 | SNP | A->G | cueO | E279G (GAG->GGG) |
| 259,748 | MOB | IS1 (+) +9 bp | phoE | coding (345-353/1056 nt) |
| 701,623 | INS | (TTGAGTTACGACCTCGTT)1->2 | nagA | coding (1129/1149 nt) |
| 2,312,001 | DEL | Δ600 bp | ompC | coding (150-749/1104 nt) |
| 2,312,023 | DEL | Δ633 bp | ompC | coding (95-727/1104 nt) |
| 4,186,532 | SNP | A->G | rpoC | K395E (AAA->GAA) |
| Strain 5 | | | | |
| 137,092 | SNP | C->T | cueO | R4C (CGT->TGT) |
| 137,093 | SNP | G->T | cueO | R4L (CGT->CTT) |
| 523,224 | SNP | C->T | ybbP, rhsD | intergenic (+394/-37) |
| 702,244 | SNP | T->G | nagA | T170P (ACC->CCC) |
| 702,276 | SNP | A->C | nagA | L159R (CTG->CGG) |
| 1,532,261 | SNP | C->A | ydcC | S149Y (TCT->TAT) |
| 2,312,671 | INS | (+) T | ompC | coding (79/1104 nt) |
| 2,312,740 | DEL | Δ1 bp | ompC | coding (10/1104 nt) |
| 3,815,811 | SNP | C->T | pyrE, rph | intergenic (-43/+52) |
| 4,183,826 | SNP | C->A | rpoB | A861E (GCG->GAG) |

Table S2: **Specific mutations from ALE.** Extended mutation information from ALE process, broken down by strain.

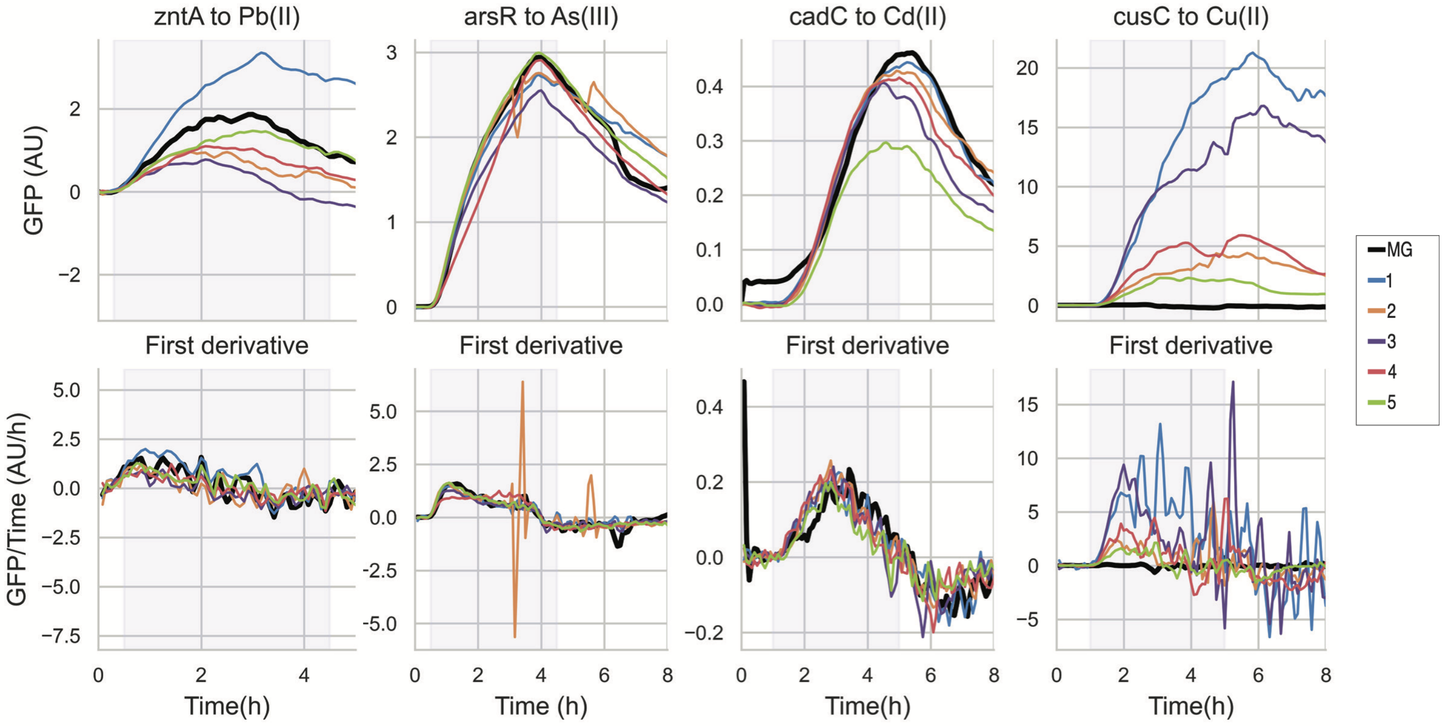


Figure S6: **Adapted strains reveal spectrum of response profiles.** Top row displays time traces of zntA, arsR, cadC, and cusC WCBs, in comparison to the MG1655 WCB. Solid lines represent mean fluorescence measured from N=8 HD traps nfor each strain. Black line represents the MG1655 WCB, while all other lines represent evolved WCBs. Heavy metal inductions were 1 ppm for all. Bottom row displays first derivative of fluorescence time traces from top row. Blue shaded region indicates the four-hour induction period.

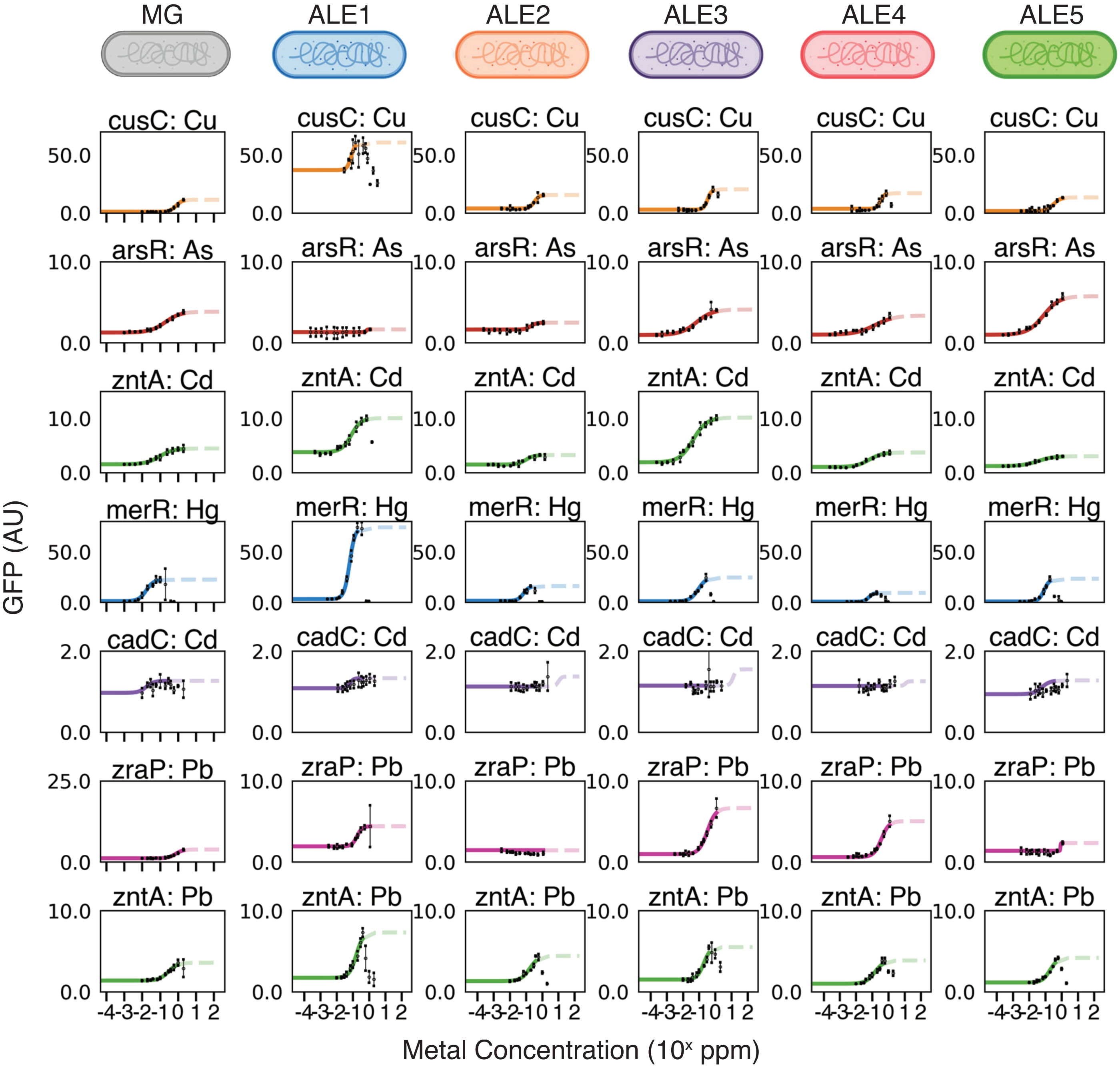


Figure S7: **Dose-response curves of evolved strains in high salinity conditions.** Batch experiment in Tecan fluorescence plate reader to compare dose-response behaviors of MG1655 WCBs to evolved WCBs in background of HM9/seawater. Concentrations indicated by dashed line affect cell growth and were not used to fit the dose-response curve. Error-bars represent standard deviation of triplicate samples.

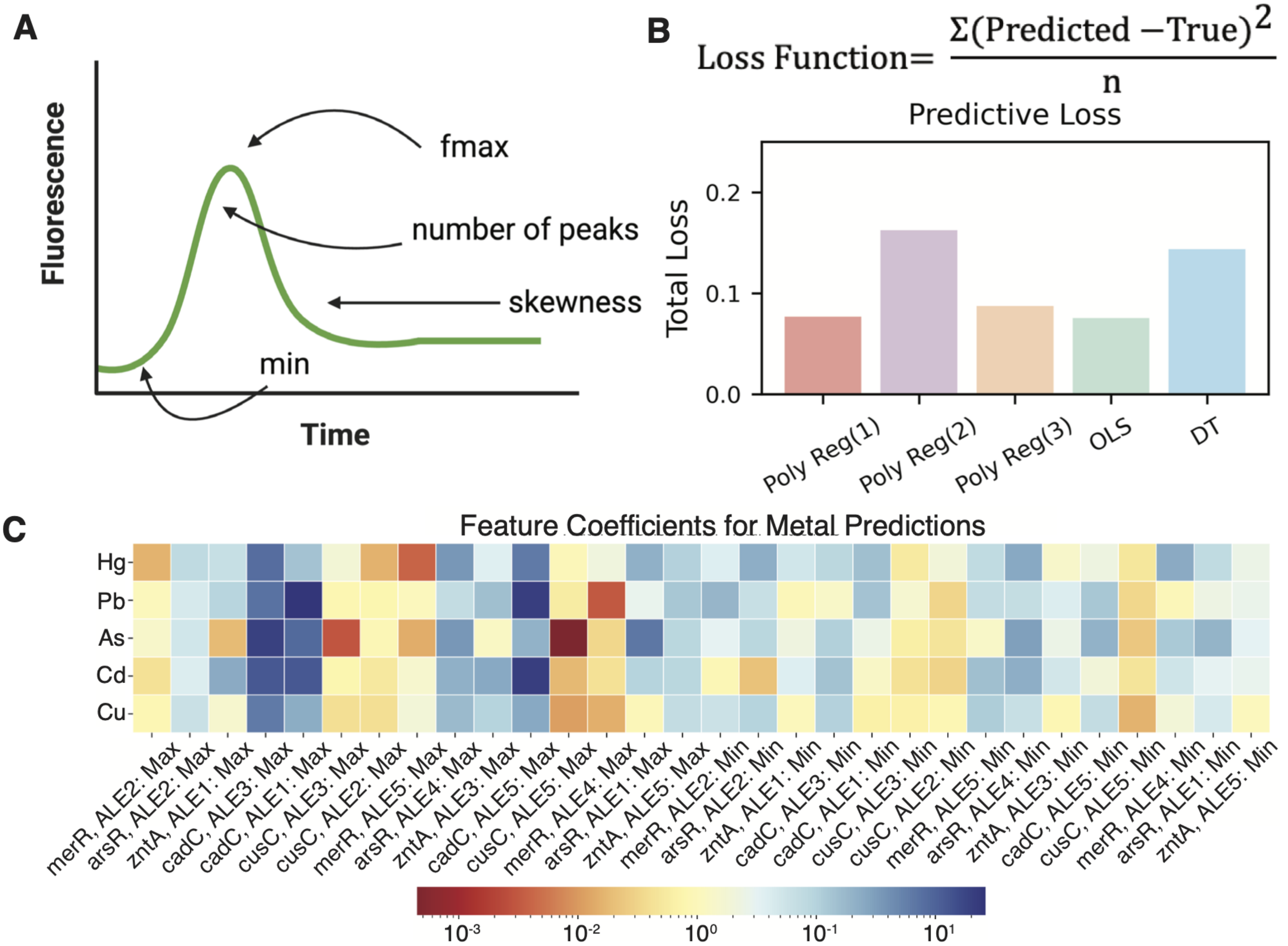


Figure S8: **Feature engineering and supervised learning model selection.** (A) The maximum, minimum, number of peaks, and skewness were among common time-series features that were evaluated. These 4 exhibited the widest distribution, indicating potential to contain information useful to classification. (B) A loss function was applied as a standard metric for evaluating predictive loss in models. Total predictive loss of first, second, and third degree polynomial regression models, ordinary least squares, and decision tree models is shown. (C) Feature coefficients for the max and min features in the ordinary least squares classification are shown to illustrate how the model makes decisions.

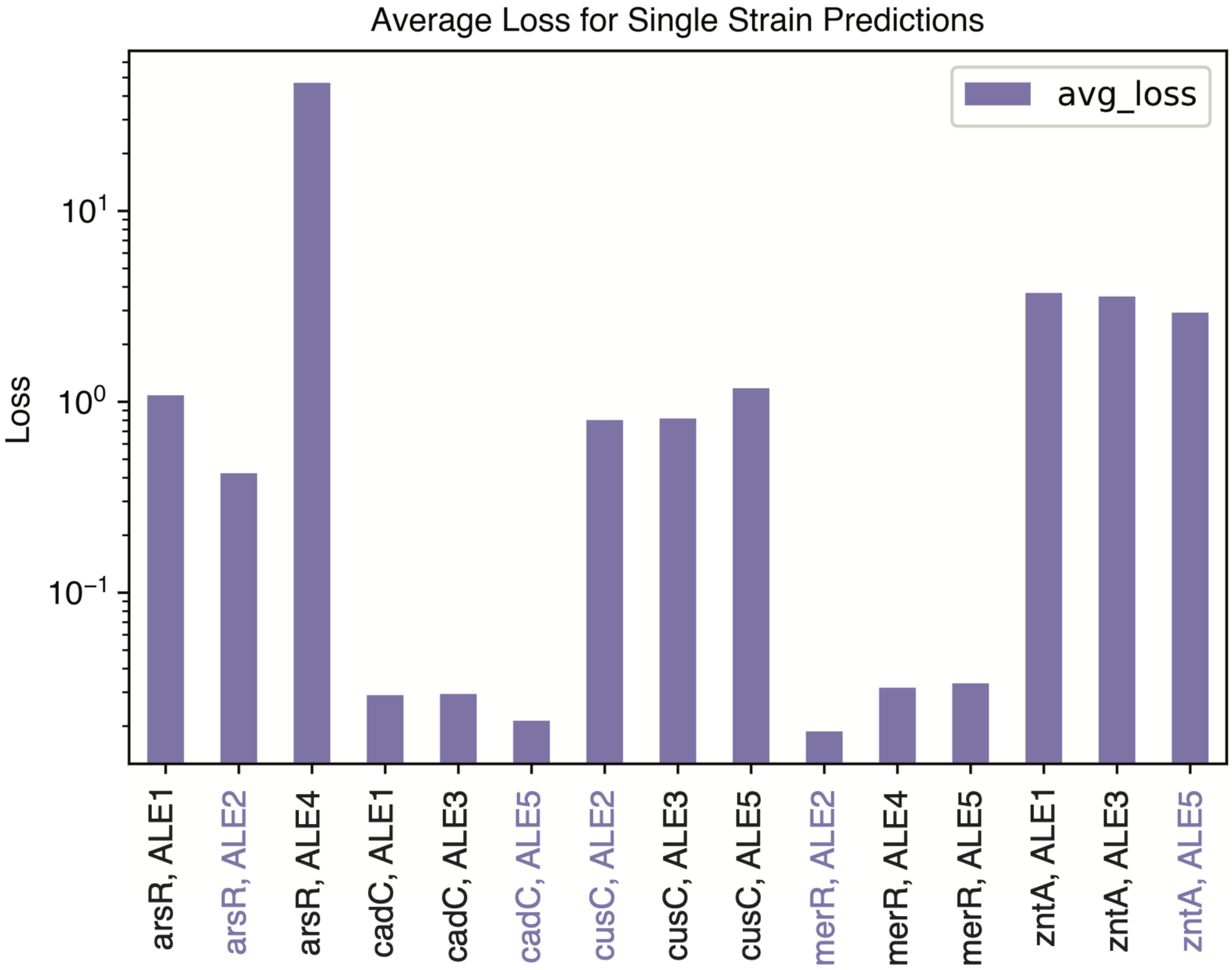


Figure S9: **Selection of single-target WCBs with optimal performance.** Each WCB's response to its intended target in each test combination was measured using its corresponding dose-response curve. The average loss of each strain across all test combinations is displayed. Each heavy metal sensor's strain exhibiting the lowest average loss was taken as the "best" single-target biosensor to compare with the consortium biosensor output.

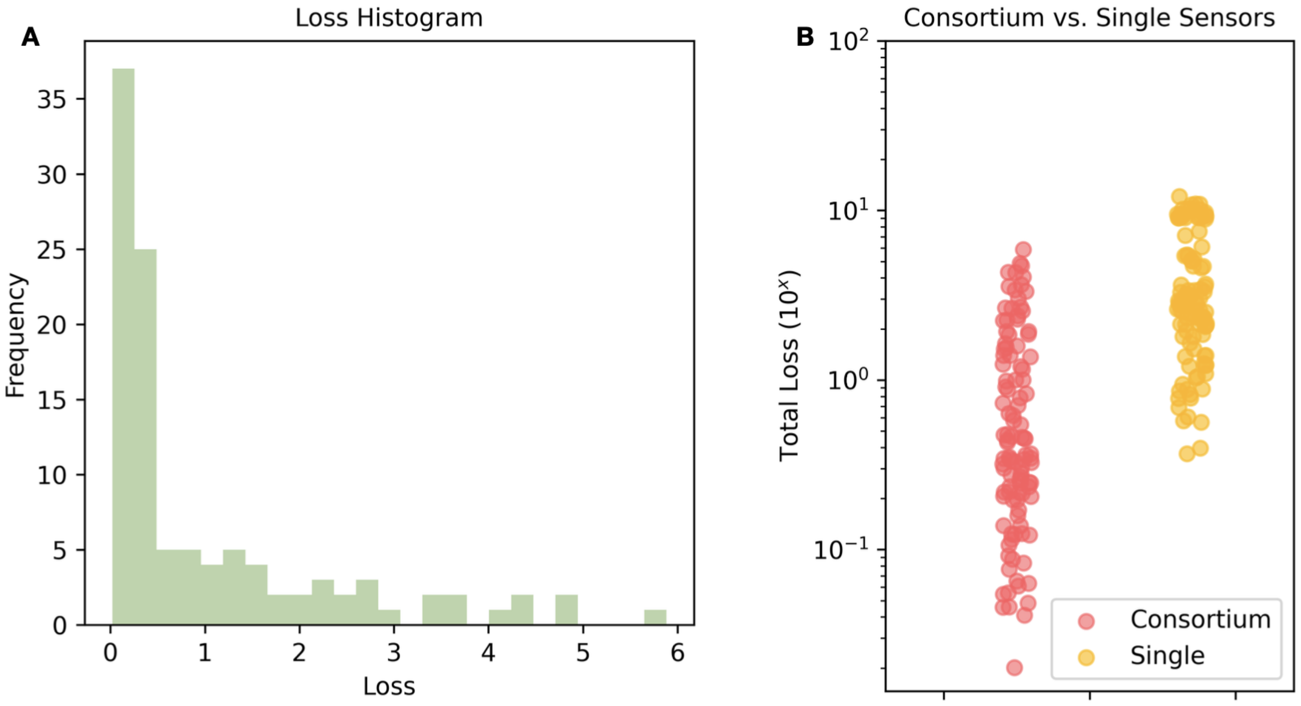


Figure S10: **Consortium performance exceeds that of single-target sensors.** (A) Loss histogram of consortium responses to test combinations. (B) Loss density plot comparing consortium and single-target biosensor paradigms.

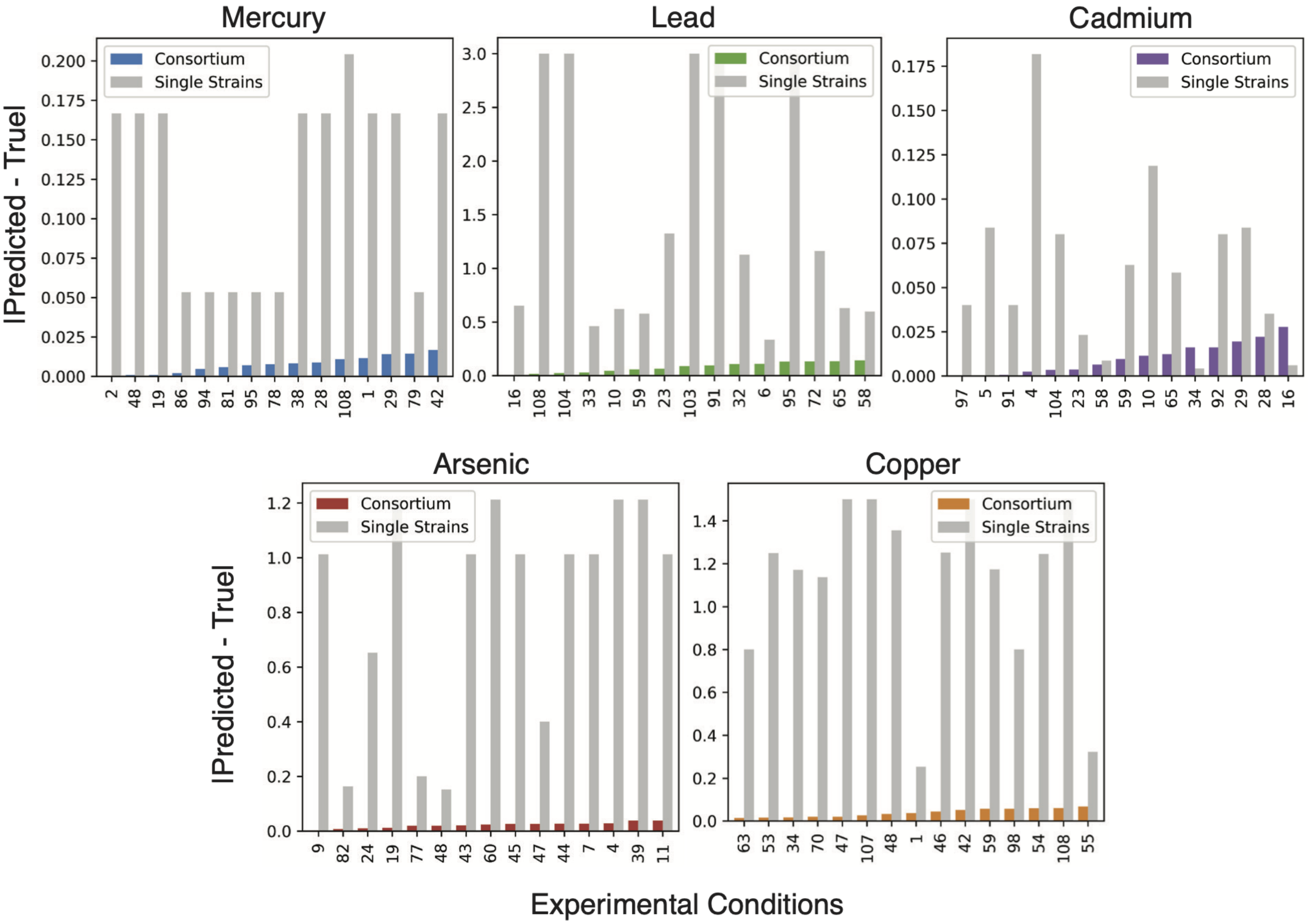


Figure S11: **Consortium v. Single-target biosensor performance, by heavy metal.** Absolute difference for the top 15 consortium predictions and their corresponding single-target predictions, organized by heavy metal. Experimental condition labels correspond to those in Figure 6C. The proportion of experimental conditions in which the consortium outperforms the single-target predictions for each heavy metal is as follows: 68% for mercury, 78% for lead, 90% for arsenic, 46% for cadmium, and 86% for copper.

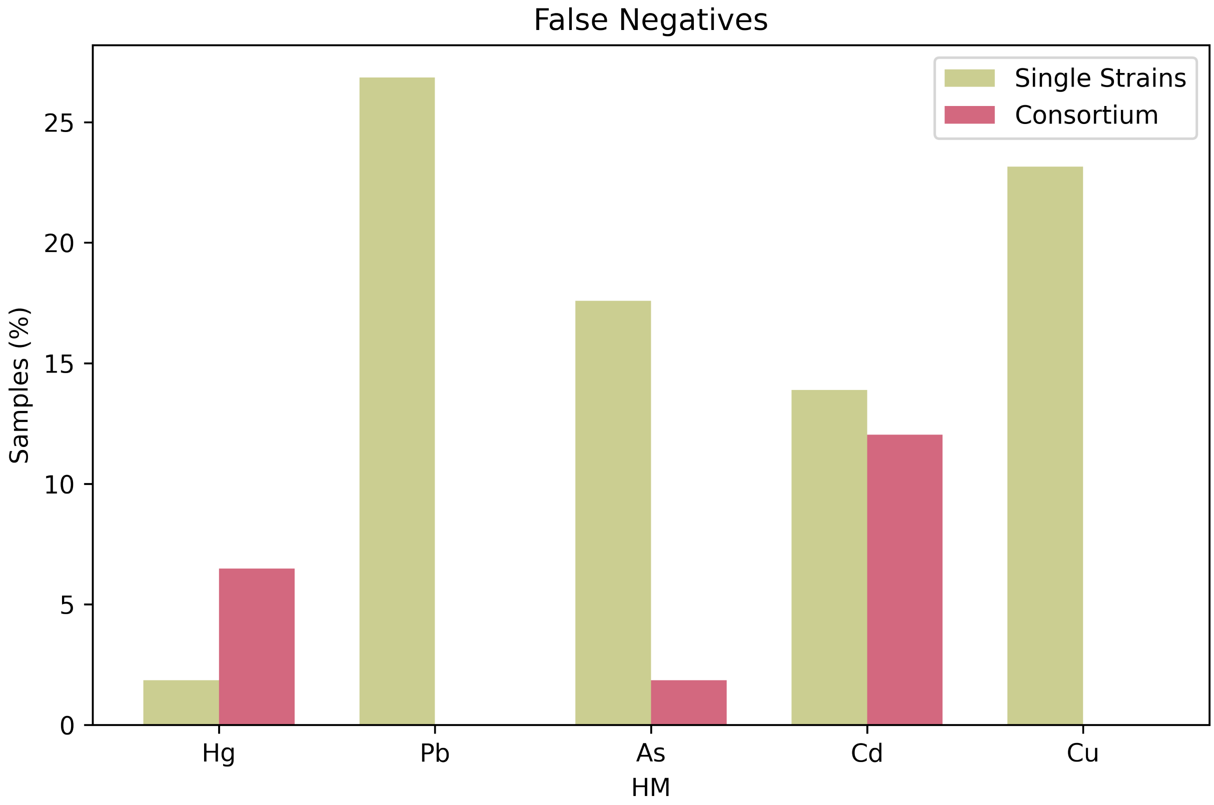


Figure S12: **Consortium v. single-target biosensor false negatives.** Count of false negatives output by consortium v. single-target biosensor predictions, by heavy metal. False negatives are defined as outputs 0 and below for consortium, and readouts below the lower limit of the biosensor's Hill function for single-target biosensors.

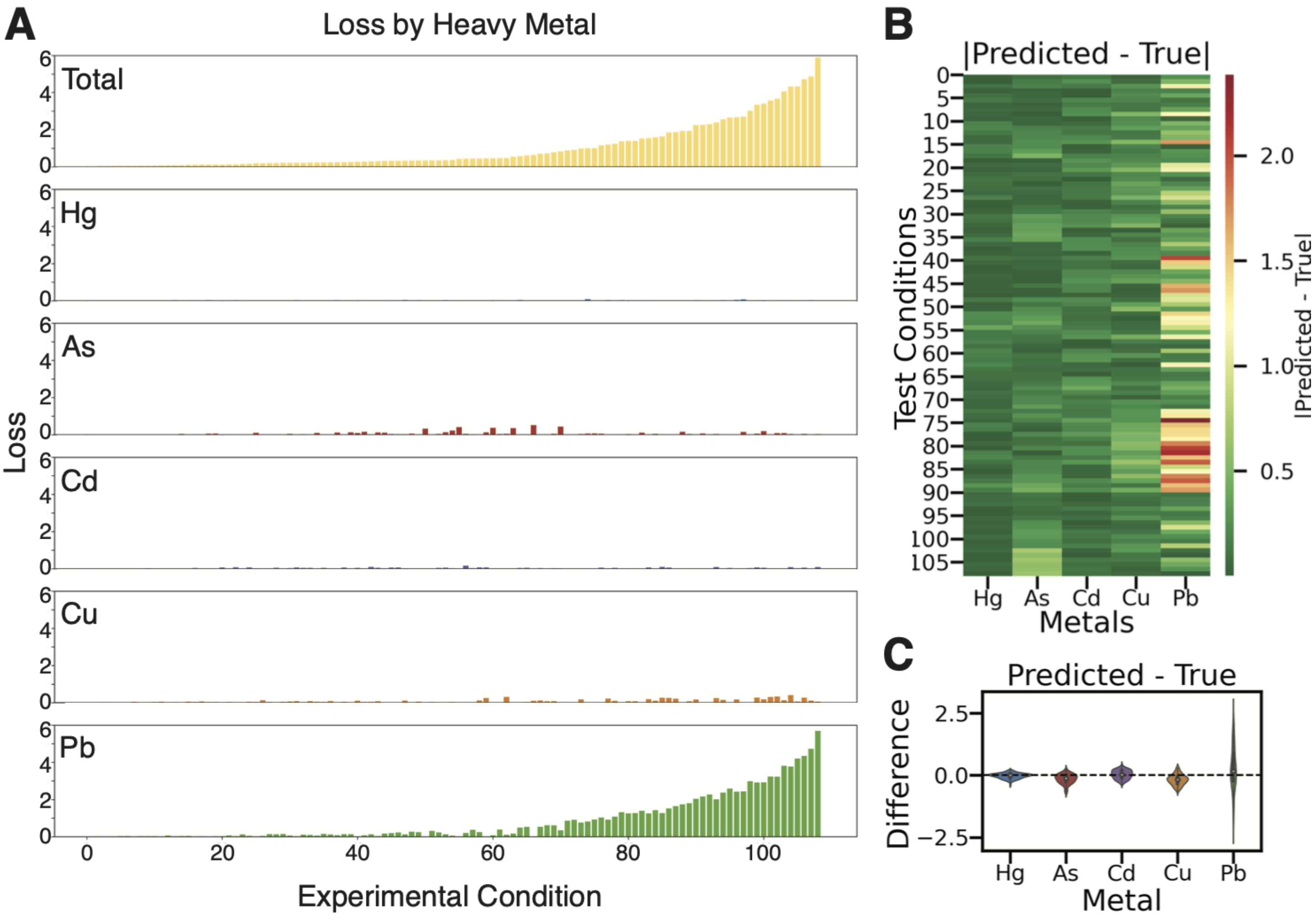


Figure S13: **Analysis of loss identifies where consortium sensor needs improvement.** (A) Total loss of consortium predictions, disaggregated into loss in each heavy meta individually. Experimental conditions are the same as in Figure 6B-C. (B) Heatmap of absolute difference between predicted and true heavy metal concentrations across experimental conditions, for each heavy metal. (C) Violin plots, by heavy metal, of difference between predicted and true values.
